# Dynamic cycles between idea generation and evaluation during creative storytelling

**DOI:** 10.1101/2024.09.04.611311

**Authors:** Xitong Liang, Mingnan Cai, Gaohan Jing, Chengming Zhang, Emily S Nichols, Li Liu

## Abstract

Many theories support the notion that creative thinking involves a dynamic transition between idea generation and evaluation, but there is limited evidence of how these two stages interact. The dual process model proposes that spontaneous thinking and deliberate thinking drive these stages and the transitions between them, but there is a debate over whether they operate in parallel or in sequence. To address these gaps, we conducted a functional magnetic resonance imaging (fMRI) study in 41 college students during a creative storytelling task. By analyzing the dynamic brain state, we link the brain states with idea generation and evaluation. The transition patterns between brain states provide evidence for dynamic circulation between idea generation and evaluation during creative storytelling. Using a deep learning approach, we demonstrate an alternating interaction between spontaneous and deliberate thinking, driving the idea generation and evaluation stages and the transitions between them. These findings deepen our understanding of the cognitive and neural mechanisms underlying creative thinking.

## 1. Introduction

*Ideas rose in crowds; I felt them collide until pairs interlocked, so to speak, making a stable combination.* Henri Poincaré, *Mathematical Creation*, 2000.

Poincaré viewed creative thinking as a complex process with various stages where countless inspirations can emerge and integrate instantly, cycling throughout the process. Psychological research has highlighted the crucial role of idea generation and evaluation in creative thinking (Runco and Acar, 2012; Girn et al., 2020; Finke et al., 1992). To explain how the two stages work together, the two-fold model of creativity proposes a cyclic motion between idea generation and evaluation (Kleinmintz et al., 2019). However, evidence to support this cycling between idea generation and evaluation remains scarce. A behavioral study of creative design used a linkograph to show that, during a time frame of creative decision making lasting approximately 7 seconds, idea generation and evaluation appear to coexist (Goldschmidt, 2016). However, it is unknown whether during this time period, the two processes are truly simultaneously active, or are oscillating quickly. Investigating neural mechanisms on a finer time scale could clarify if idea generation and evaluation cycle frequently in creative thinking (Goldschmidt., 2016; Lloyd-Cox et al., 2022).

Then, what type of thinking drives these two stages and the possible cycles between them? The dual-process model considers both spontaneous and deliberate thinking as playing important roles in creative thinking (Sloman, 1996; Evans and Stanovich, 2013). Despite no direct one-to-one match, spontaneous thinking more frequently drives idea generation, whereas deliberative thinking bolsters evaluation (Allen and Thomas, 2011; Paul et al., 2015). However, the alternating or parallel operation of the two thinking types remains controversial in dual-process theory (Paul et al., 2015). While the dual-process model has been widely accepted, consensus on the modes of these thinking types remains elusive (Chater and Schwarzlose, 2016; Evans, 2009). The parallel hypothesis, proposing simultaneous spontaneous and deliberate thinking, conflicts with cognitive economy by demanding more resources (De Neys, 2022). Conversely, the alternate hypothesis does not fully clarify the shift between spontaneous and deliberate thinking (De Neys, 2012; Evans, 2019). Given the difficulty in observing thinking types and transitions, studying neurological indicators could enhance our understanding of their interaction, potentially resolving aforementioned controversies (Paul et al., 2015).

Research on the neural basis of creative thinking focused on brain networks’ interactions, associating the default mode network with spontaneous thinking and the control network with deliberate thinking (Beaty et al., 2019; Belden et al., 2019; Vartanian et al., 2018; Zhu et al., 2017; Xie et al., 2021). Other brain networks such as the attention and visual networks have also been implicated in creative thinking (Beaty et al., 2017; Takeuchi et al., 2020; Sun et al., 2019; He et al., 2022). The dynamic framework of spontaneous thinking highlights fluctuations in large-scale networks, viewing idea generation as a spontaneous thinking process involving default mode, sensory, and attention networks. Idea evaluation involves interaction between spontaneous thinking and deliberate constraints. Deliberate constraints are related to the dynamic fluctuations of the control and attention networks. The interaction and variability between the two thinking modes facilitate stage maintenance and transitions (Christoff et al., 2016). The above framework can guide the identification of networks related to each type of thinking.

Definitive evidence of a cyclical pattern in idea generation and evaluation during creativity is limited. Further, debate persists on whether spontaneous and deliberate thinking processes occur in an alternating or parallel manner. Here, we address these issues using the advanced LEiDA method, which examines brain states and transitions on a relatively fine time scale (Deco et al., 2019; Kurtin et al., 2023), to investigate the dynamics of brain states potentially linked to idea generation and evaluation in creative storytelling. This method, utilizing data co-activation patterns, provides higher temporal resolution (1 TR) than functional connectivity methods (based on long-term time series correlation), enabling a more precise capture of brain dynamics. We then characterize the brain networks related to spontaneous and deliberate thinking based on the dynamic framework proposed by Christoff et al. (2016). Finally, utilizing deep learning techniques, we examine the interaction between these two modes of thinking in order to test the hypothesis of their alternating and parallel operation.

## 2. Materials and Methods

### 2.1 Participants

Forty-seven adult college students were recruited as participants. After excluding those with excessive head movement and program errors during the experiment, data from a total of 41 participants (18 males and 23 females) were analyzed. The mean age of the participants was 22.15 years (SD = 1.65, range = 18-25). All participants were right-handed and native Chinese speakers with normal hearing and normal or corrected-to-normal vision. Participants had no psychiatric disorders and had not taken medication that affects the nervous system. This study was approved by the Research Ethics Committee of Beijing Normal University, and all the participants provided written informed consent.

### 2.2 Behavioral tests

To evaluate intelligence and creative abilities, we conducted the following behavioral tests:

#### 2.2.1 Remote Associates Test

The Chinese version of the Remote Associates Test (RAT) assesses convergent thinking, an essential component of creative thinking ability (Wu and Chen, 2017). The test comprises 20 items, with each item presenting three Chinese characters to the participants. Participants are required to find a single Chinese character that could form a word with all three given characters. The test is timed at 20 minutes, and the score is calculated based on the number of correctly answered questions. A higher score indicates superior convergent thinking ability.

#### 2.2.2 Alternative Uses Test

The Alternative Uses Test (AUT) assesses divergent thinking ability, a crucial component of creative performance (Guilford, 1967). Participants are asked to generate as many alternative uses as possible for plastic bottles and ropes within two minutes each. The average number of uses generated for both items is calculated to determine the participant’s score for divergent thinking performance. A higher score indicates superior divergent thinking ability.

#### 2.2.3 Creativity Assessment Framework

The creativity assessment framework, from the Programme for International Student Assessment (PISA), assesses creative writing ability (Foster and Schleicher, 2022). Participants are asked to complete three creative writing prompts within a time limit of 40 minutes. The prompts are scored on a scale of 5, 10, and 10 respectively. Two graduate students studying Chinese language and literature provided independent ratings for each prompt, and we calculated the consistency coefficient between the two raters (r=0.73). The average score from both raters was taken as the participant’s final creative writing score. A higher score indicates better creative writing performance.

#### 2.2.4 Raven’s Standard Progressive Matrices

The general fluid nonverbal intelligence was assessed by the Raven’s Standard Progressive Matrices (Raven, 1938). The participants were asked to select a suitable graphic which can fill in a large graphic based on the pattern presented. The entire test contained 60 items. The total score of correct answers by a participant was then converted into a percentage level.

#### 2.2.5 Wechsler Adult Intelligence Scale-Chinese Revision (WAIS-RC)

Verbal intelligence was assessed by using the verbal dimension of the Wechsler Adult Intelligence Scale-Chinese Revision (WAIS-RC) (Wechsler, 1981). It consists of six core subtests: Vocabulary, Comprehension, Information, Similarities, Digit Span, and Arithmetic. The verbal scale scores of the subjects were obtained by adding up the scores of these six subtests, and then the standard scores of the subjects were obtained based on their scale scores.

### 2.3 fMRI experiment

The experimental design is shown in **Figure 1**. Participants were asked to perform two storytelling tasks: a creative storytelling task and an uncreative storytelling task, a design used in a previous study of creativity (Howard-Jones et al., 2005). The experiment consisted of three runs, each containing four blocks; within each run, two blocks were for the creative storytelling condition and two were for the uncreative storytelling condition. A 20-second rest period was inserted between blocks, and the conditions were alternated in each run. The total duration of each run was 388 seconds. The entire experimental task includes 24 trials (12 trials belong to creative story, other trials belong to uncreative story), totaling approximately 20 minutes.

**Figure 1.**
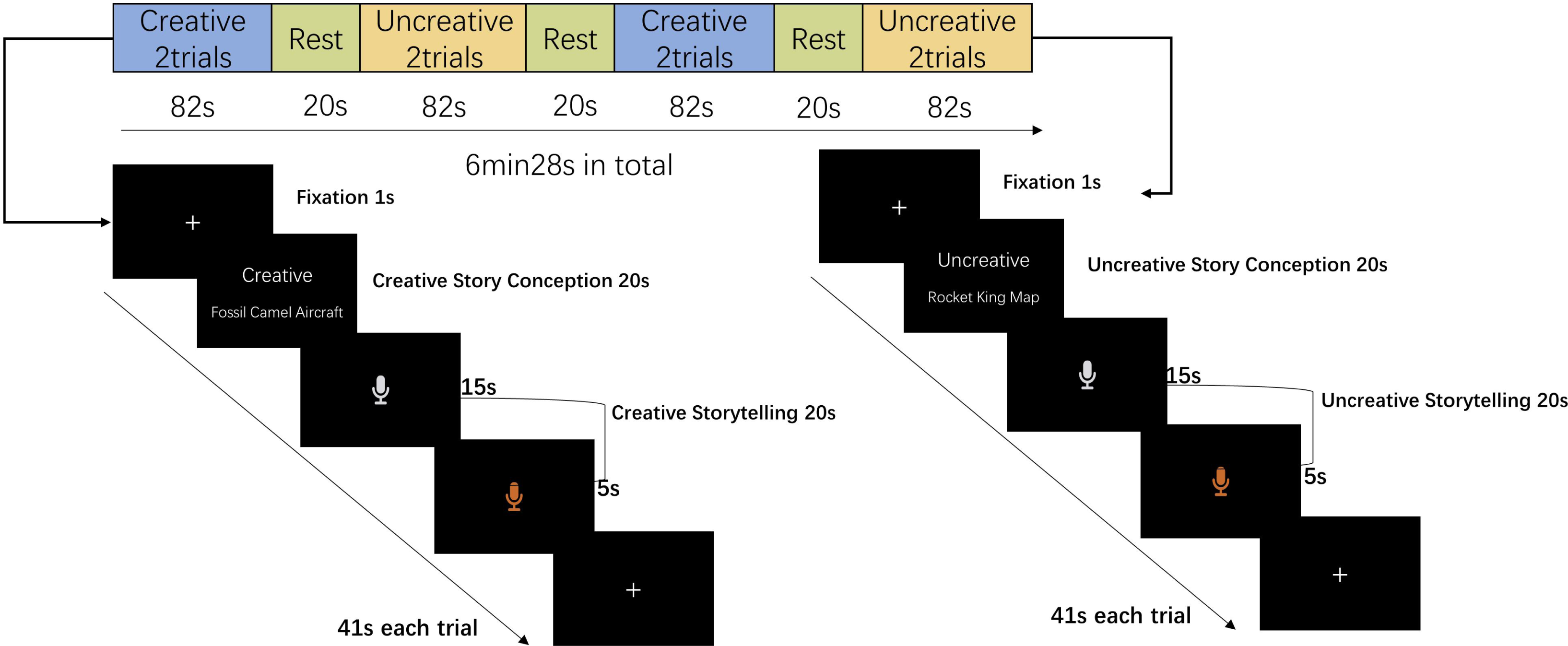
The entire experimental process.

Each block had two trials. For a trial in the creative storytelling condition, participants were shown a fixation point for approximately 1 second before being presented with task requirements (creative) and three words without obvious semantic associations. They were then given 20 seconds to produce a creative story, which was called the story conception stage. When a microphone icon appeared on the screen, participants were asked to verbally report their story within 20 seconds, which was called the storytelling stage. When the story telling stage reached 15s, the microphone sign on the screen turned red to remind the participants that there was only 5s left. The same process was repeated for the trials in the uncreative storytelling condition. However, in the conceptual stage of this condition, the task requirement presented on the screen is “uncreative”. Before the formal experiment begins, there is a practice session to ensure that participants understand the experimental requirements. These requirements include telling a creative story under creative conditions, and telling a plain, uninteresting story under uncreative conditions. To eliminate the effect of words themselves, each set of words were counterbalanced across participants. That is, each set of words was used for creative storytelling by half of the participants and for uncreative storytelling by the other half.

The creativity of each participant’s story was rated on a 5-point scale by two raters, and the consistency of the ratings was 0.73 (measured through Pearson correlation coefficient).

### 2.4 Image acquisition

Brain imaging data were collected using a Siemens 3T MRI scanner at the State Key Laboratory of Cognitive Neuroscience and Learning at Beijing Normal University. For fMRI acquisition, we used an accelerated single-shot T2 weighted gradient echo planar imaging (EPI) with parallel imaging method. The scanning parameters were as follows: pulse repetition interval (repetition time, TR) was 1000 ms; echo time (TE) was 30 ms; flip angle was 64°; slice thickness was 3 mm; number of slices was 45; FOV was 204×204 mm^2^; scanning matrix was 68×68 and voxel size was 3×3×3 mm^3^. High-resolution structural (T1) images were also acquired, with the following parameters: number of axial slices was 208; slice thickness was 1 mm; FOV was 256×256 mm²; voxel size was 1×1×1 mm^3^; pulse repetition interval (repetition time, TR) was 2530 ms; echo time (TE) was 2.27 ms.

### 2.5 Data analysis

#### 2.5.1 Image preprocessing

Raw data were labeled using the Brain Imaging Data Structure (BIDS) format and underwent preprocessing using fmriprep (Esteban et al., 2018). This process included registration, artifact correction, MNI spatial standardization with a resolution of 3×3×3, motion parameter estimation, and spatial smoothing with an isotropic, Gaussian kernel of 6mm FWHM. To eliminate head movement, we employed the ICA-AROMA algorithm from FSL (Pruim et al., 2015). Two participants were excluded due to excessive movement greater than 3mm.

We then used the latest Schaefer parcellation (Yan et al., 2023) to divide the brain into 100 areas, each corresponding to a specific label that indicated a brain region’s involvement in the yeo17 network. Each brain region was then filtered using a bandpass filter between 0.02-0.1Hz. For each participant, each of the four conditions (creative story conception, uncreative story conception, creative storytelling, and uncreative storytelling) consisted of 240 time points (12 trials x 20s, 1 TR=1s).

#### 2.5.2 Whole brain activation analysis

After data preprocessing, used a generalized linear model to conduct statistical analysis on the BOLD signal of each participant under each task condition (creative story conception, uncreative story conception, creative storytelling, uncreative storytelling, and rest conditions) (FSL6.0; Jenkinson et al., 2012). Then we calculated the contrast between the conditions of interest to obtain whole-brain activation levels. The contrasts of interest were as follows: creative conception-rest, uncreative conception-rest, creative conception-uncreative conception, creative storytelling-rest, uncreative storytelling-rest and creative storytelling-uncreative storytelling. To avoid type 1 errors, we used FSL’s cluster level correction with a threshold of Z>3.3 and p<0.001 for each contrast.

#### 2.5.3 Investigate the cyclic patterns between idea generation and evaluation

To investigate the cyclic patterns between idea generation and evaluation on a finer time scale, we initially search for neural representations for these two stages. We used a brain region synchronization method to capture the brain’s state at each TR. Brain states showing significant differences under creative versus uncreative conditions are considered as potential representations for creative thinking. By analyzing the time-varying patterns of these key states, we try to link the brain states with the two stages. Furthermore, if these states of the participants are related to their creative behavioral performance in different aspects, it may imply a relationship between brain states and these stages. Calculating the transition patterns among these states allows us to explore the cyclic nature between idea generation and evaluation. The specific analysis process is as follows:

##### 2.5.3.1 Determining the states of the brain

###### Dynamic BOLD phase-locking analysis

In line with Cabral et al. (2017), we begun by applying a Hilbert transform to the BOLD signal to derive the phase of each brain region at every time point, denoted as 𝜃 (𝑛, 𝑡). We then computed the dynamic phase-locked matrix 𝑑𝑃𝐿 (𝑛, 𝑝, 𝑡) to assess the BOLD phase consistency between any two brain regions across the entire brain at each time point *t*, utilizing the following formula:

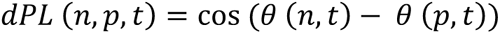

We considered the two brain regions to be more synchronized when the phase difference between their BOLD signals was less than 90 °, and assigned a positive value to the 𝑑𝑃𝐿 (𝑛, 𝑝, 𝑡). Conversely, if the phase difference exceeded 90 °, we assigned a negative value to the 𝑑𝑃𝐿 (𝑛, 𝑝, 𝑡).

Thus, for each condition, the dynamic phase-locked matrix (𝑑𝑃𝐿) for each participant was a three-dimensional tensor with dimensions N x N x T, where N equals 100 and represents the number of brain regions, and T equals 240 and denotes the total number of time points.

###### Leading Eigenvector Dynamics Analysis (LEiDA)

The Leading Eigenvector Dynamics Analysis (LEiDA) method, as proposed by Cabral et al. (2017), can be used to reduce the dimensionality of dynamic phase-locked matrices (𝑑𝑃𝐿). This technique involves using the leading eigenvector 𝑉1(𝑡) of each 𝑑𝑃𝐿 (𝑡) to represent the primary phase orientation of the matrix. The symbols of N elements in 𝑉1(𝑡) are employed to depict the phase relationship between brain regions. When all elements share the same symbol, it signifies that all BOLD signals are aligned in the same direction, indicating global synchronization. If an element possesses both positive and negative values, the brain region is divided into two communities based on these polarities. As 𝑉 and −𝑉 represent identical states, a convention is adopted to ensure that the majority of elements are negative. This implies that a smaller subset of brain regions is synchronized at a specific time point.

###### Detection of BOLD phase-locking state

We applied k-means clustering to the 𝑉1(𝑡) signals of all participants at each time point under each condition to identify the brain’s BOLD phase-locking pattern. The number of V1 signals obtained under each condition was 41 x 240=9840. As there is no consensus on the optimal number of k-means clusters, we followed previous research and set the range of clusters to 5-10 (Lord et al., 2019).

For each clustering result, we used the centroid of the cluster to represent the BOLD phase-locking pattern of each state. Through clustering algorithms, we were able to obtain the BOLD phase-locking state of each participant at each time point. To aid in understanding, we have created a schematic diagram illustrating the steps mentioned above, which is presented in **Figure 2**.

**Figure 2.**
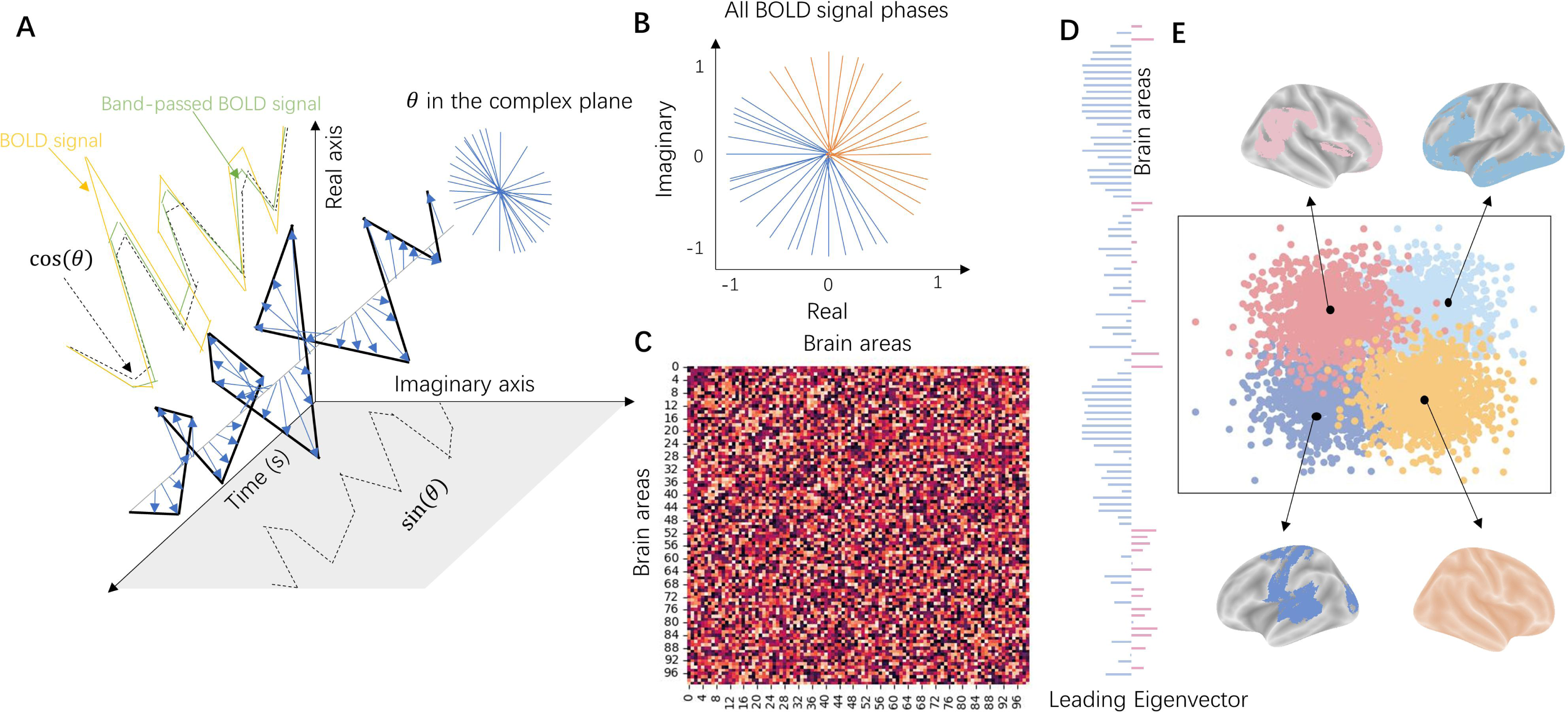
Schematic diagram of the Leading Eigenvector Dynamics Analysis (LEiDA) method. **A)** displayed the process of conducting Hilbert transform to the BOLD signal to derive the phase of each brain region at every time point. The yellow line represented the BOLD signal. The green line represented the band-passed BOLD signal. The black line represented the phase dynamics of the analyzed signal, where the real part was represented by cos (θ) and the imaginary part was represented by sin (θ). The blue arrow indicated the BOLD phase of each TR. **B)** indicated the phase of all brain regions at a certain timepoint. **C)** showed the phase-locked matrix at a certain timepoint. **D)** displayed a leading eigenvector V at a certain timepoint. In **E)**, we performed cluster analysis on the leading eigenvector V of all participants at all time points. Every colored dot presented a leading eigenvector V, and the color represented the cluster it belongs to. The black dots represented the center of each cluster. The brain map represented the synchronous state of the brain at each cluster’s center. This figure is only for the convenience of readers to understand, and the method is from Deco et al. (2019).

To search for key states in creative story conception or storytelling conditions, we defined the probability of each state occurring as the ratio of the number of time points at which each state occurs to the total number of time points. We then performed a paired t-test based on permutation between the creative story conception and uncreative story conception conditions, as well as between the creative storytelling and uncreative storytelling conditions. If the probability of a state occurrence significantly increased in creative conditions compared with uncreative conditions, it was considered as a key state. To ensure the robustness of our results, we used a Bonferroni corrected p-value threshold of α= 0.05/k (where k is the number of K-means clusters).

##### 2.5.3.2 The dynamics of brain states

###### The dwell life time of key states

In order to evaluate the duration of maintenance of key states, we calculated the dwell life time (average duration of time spent maintaining a particular state) of key states for each participant by dividing the total duration of their appearance by the total number of times they appeared. We then conducted a paired t-test to compare the dwell life time of these states between the creative and uncreative conditions.

###### The number of transitions

To quantify the extent of brain state changes during the task, we calculated the number of transitions for each participant. A transition was recorded when the state at time t+1 differed from the state at time t. We then conducted a paired t-test to compare the number of transitions between the creative and uncreative conditions. In addition, we also compared the number of transitions of key states between the creative and uncreative conditions.

###### The transition pattern between key states

To compare the differences in probability of state transition between creative and uncreative conditions for story conception and storytelling stages respectively, we first calculated the probability of each participant’s transition between key states and other states under four conditions (creative conception, uncreative conception, creative storytelling, uncreative storytelling). Specifically, we divided the number of transitions between target states by the total number of transitions. Then paired t- test was applied to each transition probability under the conditions of creative conception and uncreative conception, and the creative storytelling and uncreative storytelling conditions. We used Bonferroni correction for multiple comparison correction (p-value threshold of α= 0.05/30. For a key state, a t-test will be conducted on the bidirectional transition probability between it and the other 5 states under creative conditions compared to uncreative conditions. The number of t-tests is 5 x 2=10, and there are a total of 3 key states in the conception and storytelling stages. Therefore, the total number of t-tests is 30).

###### The probability of key states occurring over time

In order to measure the temporal variation pattern of key states, we calculated the cumulative probability change of key states for each participant during each 20- second task time window under creative/uncreative story conception or storytelling. We used 𝑛 to represent the total number of key states appearing from the 1st second onwards, and 𝑡 to represent the current time. The formula for calculation is as follows:

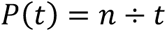

If the probability of a key state increases rapidly over time, then its cumulative probability acceleration should be greater than zero. Conversely, if it decreases rapidly, then its acceleration should be less than zero. To quantify this, we performed quadratic curve fitting on each 𝑃(𝑡). The resulting quadratic coefficient served as a measure of acceleration. We then conducted a t-test to determine if the acceleration was significantly different from zero.

##### 2.5.3.3 Linking brain states and creative behavior performance

To investigate the relationship between brain states and creative performance, we conducted a regression analysis using each participant’s RAT score, AUT score, creative writing score, and creative storytelling score as dependent variables respectively. We used the difference in dwell time of key states between creative and uncreative story conception as independent variables. Additionally, we included verbal IQ, non-verbal IQ, age, and sex as control variables in our analysis.

Before conducting the regression analysis, we first conducted a correlation analysis to select independent variables and control variables, and only those variables showing a significant correlation with the dependent variable were selected to enter the regression models. Then, we tested the overall fit of the models as well as the coefficients of the independent variables.

#### 2.5.4 Examine the parallel and alternating hypotheses of the dual process model

To examine the parallel and alternating hypotheses of the dual process model, we first identified brain networks associated with spontaneous and deliberate thinking. We then input data from these networks into parallel or alternating deep learning models to predict dynamic brain state changes. In recent years, numerous studies have utilized deep learning to simulate neural signals, aiming to comprehend patterns of neural activity (Schrimpf et al., 2021; Henningsen-Schomers et al., 2023). The attention mechanism, a deep learning method, assigns varying weighted parameters to inputs based on their significance. This mimics the human brain’s cognitive pattern of allocating limited attention to key aspects of things as needed. Notably, parallel and alternating co-attention models consider different information collaboration methods (Lu et al., 2016; Xiong et al., 2016). Therefore, comparing the performance of these models can partially explain patterns of brain network interaction, thereby testing the relevant hypotheses.

To start with, we sought to determine which brain networks were associated with spontaneous and deliberate thinking. As reported by Christoff and colleagues (2016), variations in the default mode network, attention network and sensory network are related to spontaneous thinking, while deliberate constraints on spontaneous thinking manifest in the interaction of default mode network, attention network and control network. The dynamic fluctuations and interactions between two types of thinking facilitate the dwelling and transitioning between different processing stages (Christoff et al., 2016). Consequently, we identified specific brain networks whose activity fluctuations correlate with these dwellings and transitions in key brain states. We further categorized these networks to determine their association with particular types of thinking.

Specifically, we conducted the following calculations to investigate the relationship between the dwellings and transitions of key brain states and brain networks. As the creative conception stage best reflects the variations in creative thinking, we only selected this stage for calculation. That is, during creative conception, the story must be generated, requiring creative thinking, while the storytelling stage simply requires repeating the story that has been generated. Thus, we identified key states and transitions between them during the creative story conception stage. We labeled the time points spent in key states as dwell times and those that underwent transitions as transition times. To analyze brain activity during these events, we divided the brain into 8 networks based on the Yeo17 atlas, including the visual network, somatomotor network, dorsal attention network, ventral attention/salience network, default mode network, control network, superior temporal network, and limbic network. For each network, we calculated the variation of the activation per second. We then compared the variations of the activation per second during vs. outside of the key states we have identified through the above analysis. We also compared the variations of the activation per second in these transition time points between key states with those that do not involve transitions between these key states.

Next, we classified the identified brain networks into those associated with spontaneous and deliberate thinking based on the dynamic framework (Christoff et al., 2016), and employed deep learning methods to explore the mechanisms underlying spontaneous and deliberate thinking. Co-attention mechanisms, a form of deep learning model, can be employed to handle multiple information sources, such as jointly predicting an answer based on a question and an image (Lu et al., 2016; Xiong et al., 2016). Two types of co-attention mechanisms have been proposed: Parallel and Alternating, each with distinct methods of combining information (Lu et al., 2016; Xiong et al., 2016). Both co-attention models produce attention from various information sources. However, Parallel Co-Attention model consider information from all sources simultaneously, whereas Alternating Co-Attention model process information sequentially. Before conducting model training, we first divided the data into training and testing dataset.

The labels of the data included dwell time and transition time, and the features included the activity of brain networks that significantly correlated with dwell time and transition time. The data were randomly divided into 70% for the training dataset and 30% for the testing dataset. We began by employing the Alternating Co-Attention mechanism, which alternately generates attention from different feature sources (Lu et al., 2016; Xiong et al., 2016). The specific algorithm was as follows:

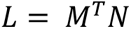

Where we combined the information of two feature matrices (i.e., multiplication) 𝑀 and 𝑁.

By calculating Softmax for the combined matrix 𝐿 row wise and column wise, we could obtain the Attention matrix for the two features:

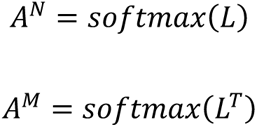

Next, we applied Attention to the feature matrix 𝑁:

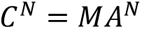

and to the feature matrix 𝑀, i.e 𝑁𝐴^𝑀^. We replaced the feature matrix 𝑀 with the feature matrix 𝑁 after adding Attention to 𝑀, that was 𝐶^𝑁^𝐴^𝑀^. As both of these steps need to be multiplied by the matrix, we could perform parallel calculations:

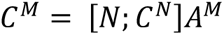

We considered it as a combination of feature information after introducing the Co-Attention mechanism, and finally we fused the information of feature 𝑀:

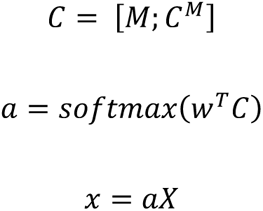

Where 𝑎 was the attention weight of feature, 𝑤 was parameters, 𝑥 was attention operation, and 𝑋 was the feature.

We next employed the Parallel Co-Attention mechanism, which generates attention from different feature sources simultaneously (Lu et al., 2016). The specific algorithm was as follows:

The first step was the same:

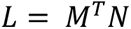

Next, we generated attention simultaneously

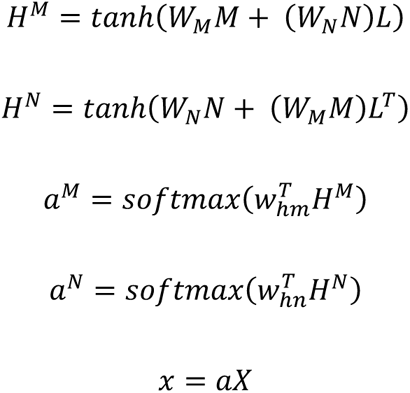

where 𝑊_𝑀_, 𝑊_𝑁_, 𝑤_ℎ𝑚_, 𝑤_ℎ𝑛_ were the weight parameters, 𝑎 was the attention weight of feature, 𝑥 was attention operation, and 𝑋 was the features. **Figure 3** shows a schematic diagram of the algorithm for easier understanding.

**Figure 3.**
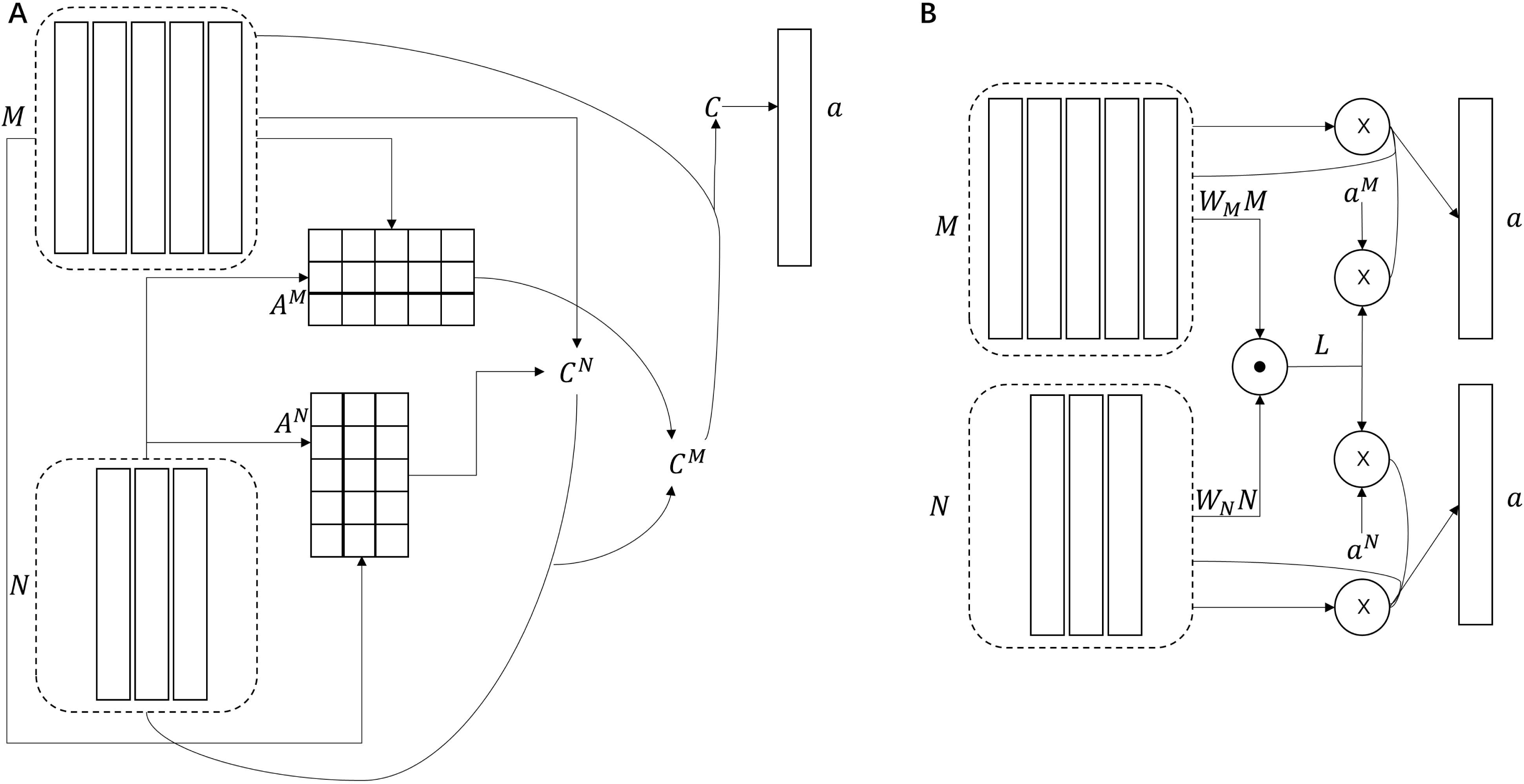
The schematic diagram of two co-attention algorithms. A) displayed the processing of Alternating Co-Attention model. B) displayed the processing of Parallel Co-Attention model. This figure is only for the convenience of readers to understand, and the method is from Lu et al. (2016) and Xiong et al. (2016).

In both mechanisms, we utilized a single-layer perceptron for the output. We employed stochastic gradient descent (SGD) to train the two models, with a learning rate of 0.01 and binary cross entropy between the target and output as the loss function. The hidden layers were set to 2, 8, 16, 24, 32, 64, 128, and 256, respectively. An epoch, which represents one complete traversal of the training data, significantly impacts the model’s generalization ability. Therefore, we varied the number of epochs between 20 and 30. We used these two models to predict the timepoints dwell in key brain states and transition between key brain states, and compared the predictive accuracy of the two models under above training parameters.

## 3. Results

### 3.1 Behavioral results

**Table 1** provides an overview of the participants’ behavioral performance. Our findings revealed a significant difference in creativity between stories told under creative and uncreative storytelling conditions (t=13.01, p<0.0001).

**Table 1.** Descriptive statistics on behavioral performance of participants.

| Variables | Mean | SD | Max | Min |
| --- | --- | --- | --- | --- |
| RAT | 13.51 | 4.12 | 21 | 2 |
| AUT | 15.15 | 4 | 29 | 8 |
| Creative Writing | 10.70 | 2.24 | 15.5 | 7 |
| Creative Storytelling (in scan) | 2.44 | 0.08 | 3.04 | 1.5 |
| Uncreative Storytelling (in scan) | 1.98 | 0.05 | 2.42 | 1.25 |
| Raven' s matrices score (non-verbal IQ) | 87.44 | 15.33 | 95 | 50 |
| WAIS score (verbal IQ) | 129.51 | 5.28 | 139 | 119 |

Notes: RAT = Remote Associates Test; AUT = Alternative Uses Test

**Figure S1** illustrates an example of the stories told by participants, demonstrating that stories created under uncreative conditions are more mundane, while those under creative conditions are more imaginative.

Besides, we performed a textbook analysis of the stories told by the participants. T-tests on the textual features of the stories under creative and uncreative conditions revealed that the mean Chinese character frequency was significantly lower in creative stories compared to uncreative ones (t=-2.13, p=0.03). The mean sentence length in creative stories was significantly higher than in uncreative stories (t= 2.41, p=0.02).

### 3.2 Results of whole brain activation analysis

Compared to uncreative story conception, creative story conception showed greater activation in extensive brain areas, including occipital cortex, precentral gyrus, precuneus gyrus, and prefrontal cortex. Similarly, compared to the uncreative storytelling, creative storytelling showed greater activation in brain regions similar to those under conception stage. **Table 2** and **Figure 4** display brain regions with significant differences between creative and uncreative conditions. Results showing creative versus rest and uncreative versus rest are presented in **Table S1** and **Figure S2**.

**Figure 4.**
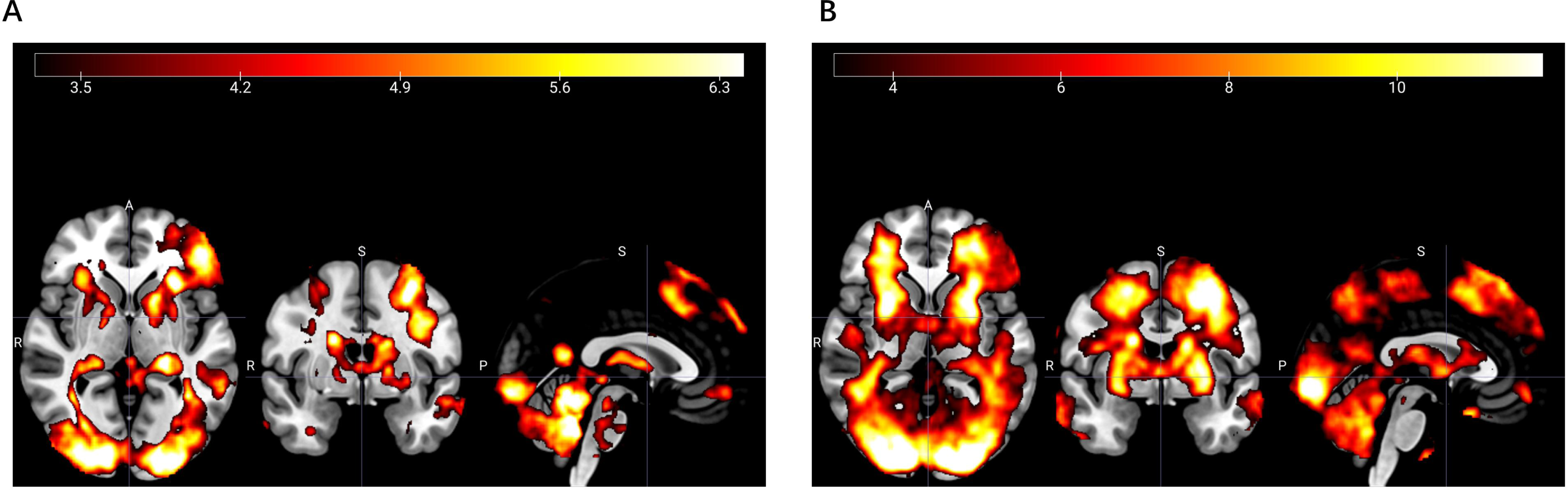
Whole brain activation under creative conditions compared to uncreative conditions. **A)** showed brain activation differences between creative and uncreative story conceptions. **B)** showed brain activation differences between creative and uncreative storytelling.

**Table 2.** Whole brain activation results under different contrasts.

| Brain regions | Volume | x | y | z | Peak<br>z |
| --- | --- | --- | --- | --- | --- |
| **Creative conception > Uncreative conception** |  |  |  |  |  |
| Right Calcarine | 541174 | 17 | -92 | -4 | 8.96 |
| Right Precentral Gyrus | 11136 | 31 | -21 | 58 | 5.59 |
| Right Paracentral Lobule | 1530 | 8 | -29 | 61 | 4.77 |
| Left Precuneus | 1366 | -6 | -40 | 62 | 4.97 |
| Left Middle Frontal Gyrus | 1291 | -2 | 51 | -12 | 5.23 |
| **Creative storytelling > Uncreative storytelling** |  |  |  |  |  |
| Right Calcarine | 220717 | 17 | -92 | -4 | 8.96 |
| Left Inferior Frontal Gyrus | 74895 | -40 | 32 | -12 | 7.92 |
| Right Calcarine | 3348 | 18 | 3 | 23 | 6.98 |
| Left Middle Frontal Gyrus | 2237 | -28 | 52 | 6 | 6.08 |
| Right Inferior Frontal<br>Gyrus | 1810 | 51 | 28 | 24 | 5.6 |
| Left Middle Temporal<br>Gyrus | 1407 | -55 | -4 | -16 | 6.04 |
| Right Insula | 944 | 32 | 26 | 2 | 5.88 |
| Right Precentral Gyrus | 861 | 43 | -30 | 54 | 5.36 |
| Right Inferior Frontal<br>Gyrus | 767 | 40 | 39 | -11 | 5.93 |

#### 3.3.1 Results related to the cyclic patterns between idea generation and evaluation

##### 3.3.1.1 The key states under creative story conception and storytelling conditions

All the variances explained by the leading eigenvector of 𝑑𝑃𝐿 matrices are greater than 50%, indicating that the leading eigenvector could capture most of the matrix’s features. Regardless of the number of clusters used in the clustering analysis, we found that during creative story conception, two states (named State 2 and State 3 when the cluster number was set to 6) had a significantly higher probability of occurrence than under uncreative story conception. The statistical significance of these findings is shown in **Figure 5A** and **Figure 5B**. Additionally, we found that a state (named State 6 when the cluster number was set to 6) had a significantly higher probability of occurrence during creative storytelling compared to uncreative storytelling. The statistical significance of this finding is shown in **Figure 5C**.

**Figure 5.**
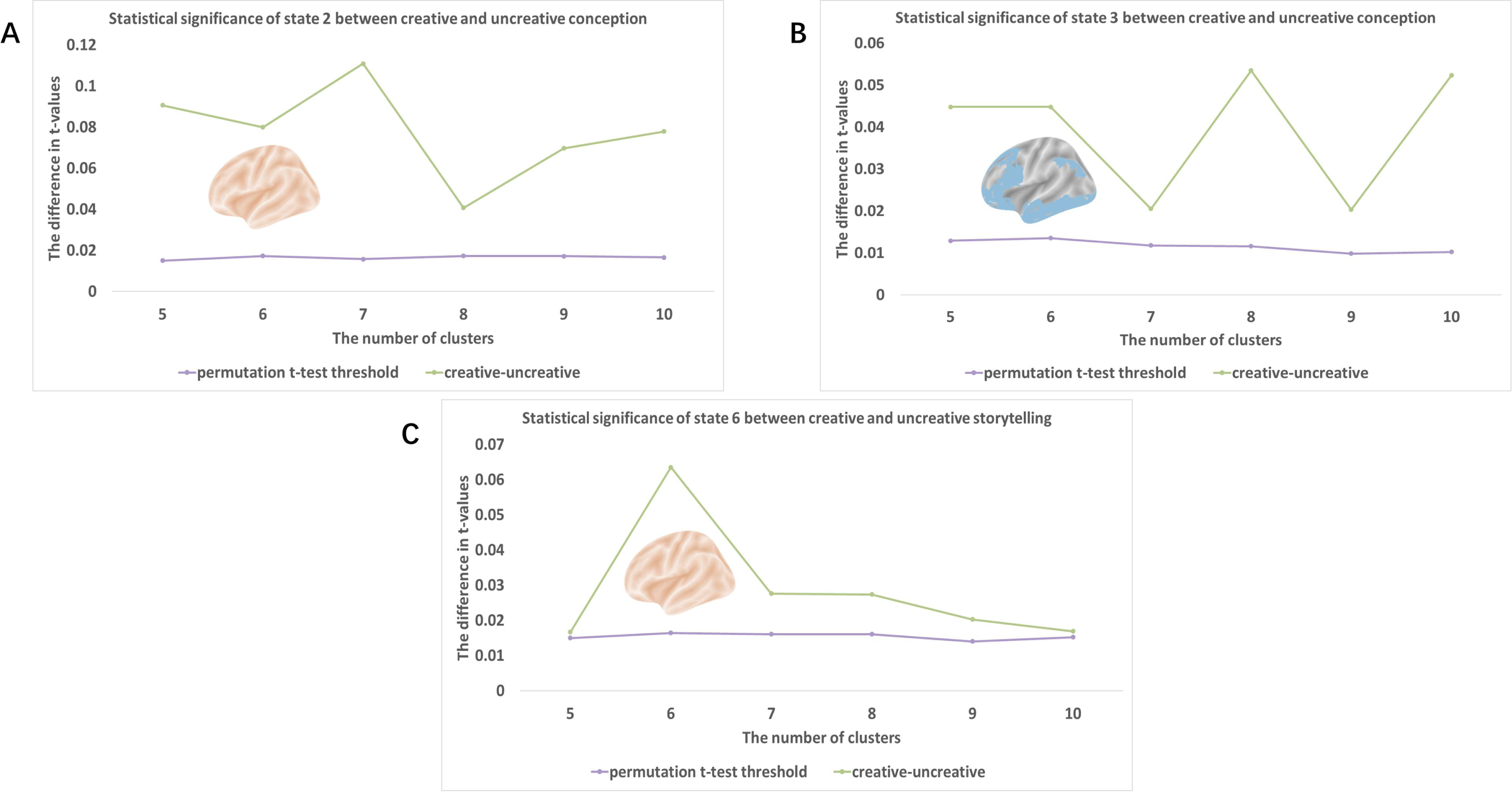
The statistical test results for state2 **(A)**, state3 **(B)** and state6 **(C)** are displayed when the number of clusters is between 5-10. The green dots and connecting lines represent the difference in t-values between creative and uncreative stage. On the other hand, the purple dots and connected lines represent the threshold for the difference in t-values when using a permutation t-test with a significance level of α= 0.05/k, where k is the number of K-means clusters. The brain maps showed the states with 6 clusters.

To ensure the robustness of our results, we performed Kendall’s W test on the synchronization pattern of brain regions in state 3 for all cluster numbers. The Kendall’s Wt was 0.94, and the p value was 3.57×10^-65^, indicating strong consistency under different numbers of clusters. State 2 and State 6 were also highly consistent across all numbers of clusters used in the clustering analysis.

We further analyzed the time points at which these key states appeared by concatenating the states of all participants under the creative story conception condition into a column according to their subject numbers. We marked the time point at which State 2 appeared as 1, the time point at which State 3 appeared as 2, and other states as 0. We conducted a Kendall W-test on this column for all cluster numbers, and the results showed that the Kendall’s Wt was 0.672 (p<0.0001), suggesting strong consistency in the time points at which these key states appeared under different numbers of clusters.

This indicates that the brain regions in key states and their timing are consistent regardless of cluster number, which serves as the prerequisite for our subsequent analysis using a specific number of clusters. We selected results with a cluster number of 6 for further analysis to demonstrate these key states. There were relatively large differences in the probability of these key states appearing between the creative and uncreative conditions, which were evident during both the conception and storytelling stages. The synchronization pattern of the brain networks for all states with a cluster number of 6 was shown in **Figure 6 and Table 3**. Among those key states, State 2 and State 6 both exhibited whole brain synchronization, while State 3 manifested as synchronization of the Control Network, Default mode Network, Visual Network, and Limbic Network (hereafter referred to as “CDVL network”).

**Figure 6.**
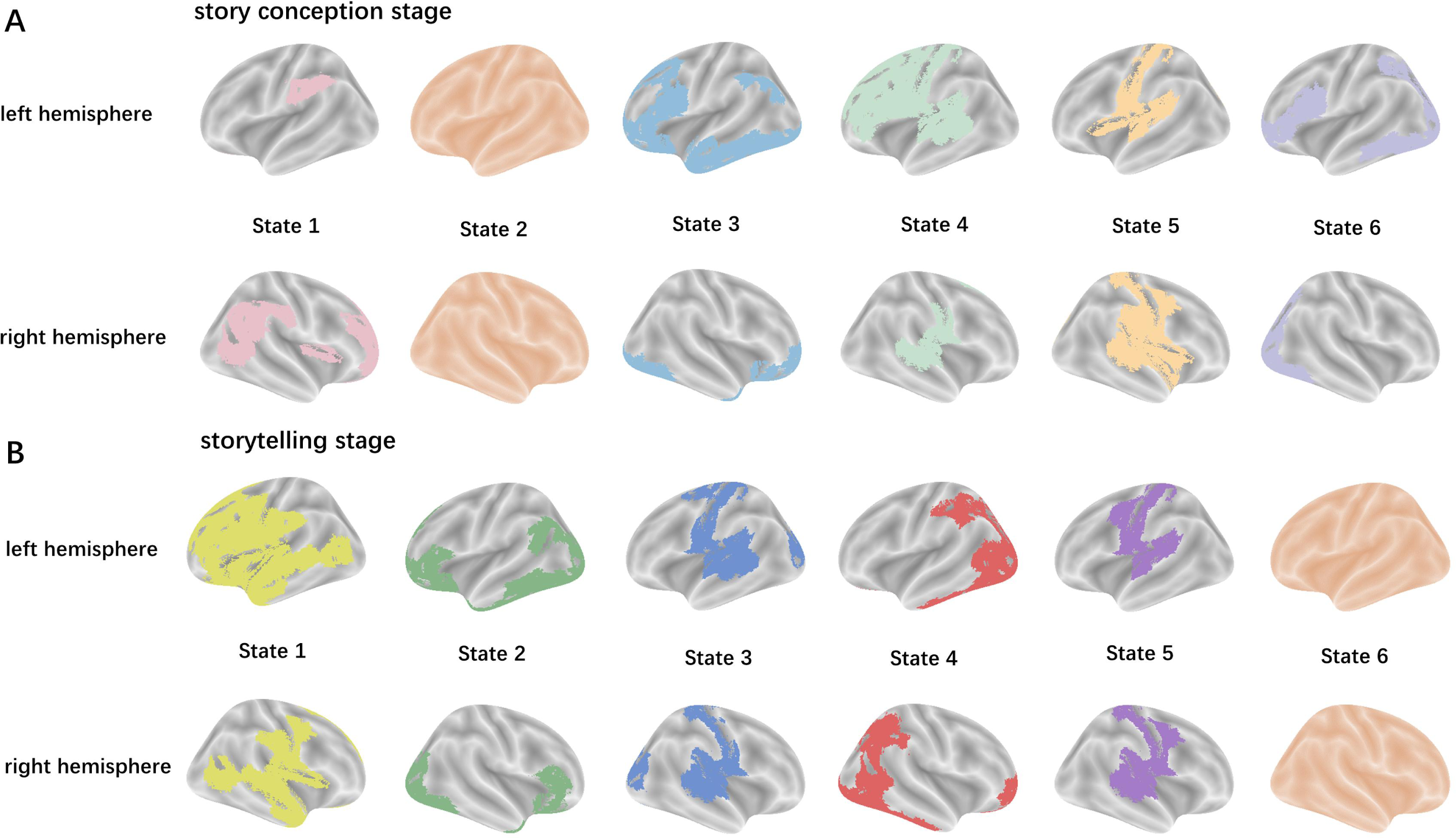
The brain synchronization patterns of 6 states during the story conception period **(A)** and storytelling period **(B)** are presented from in sequence left to right. The colored areas on the brain map represented the synchronized brain regions in this state. Different colors were used to distinguish different states.

**Table 3.**
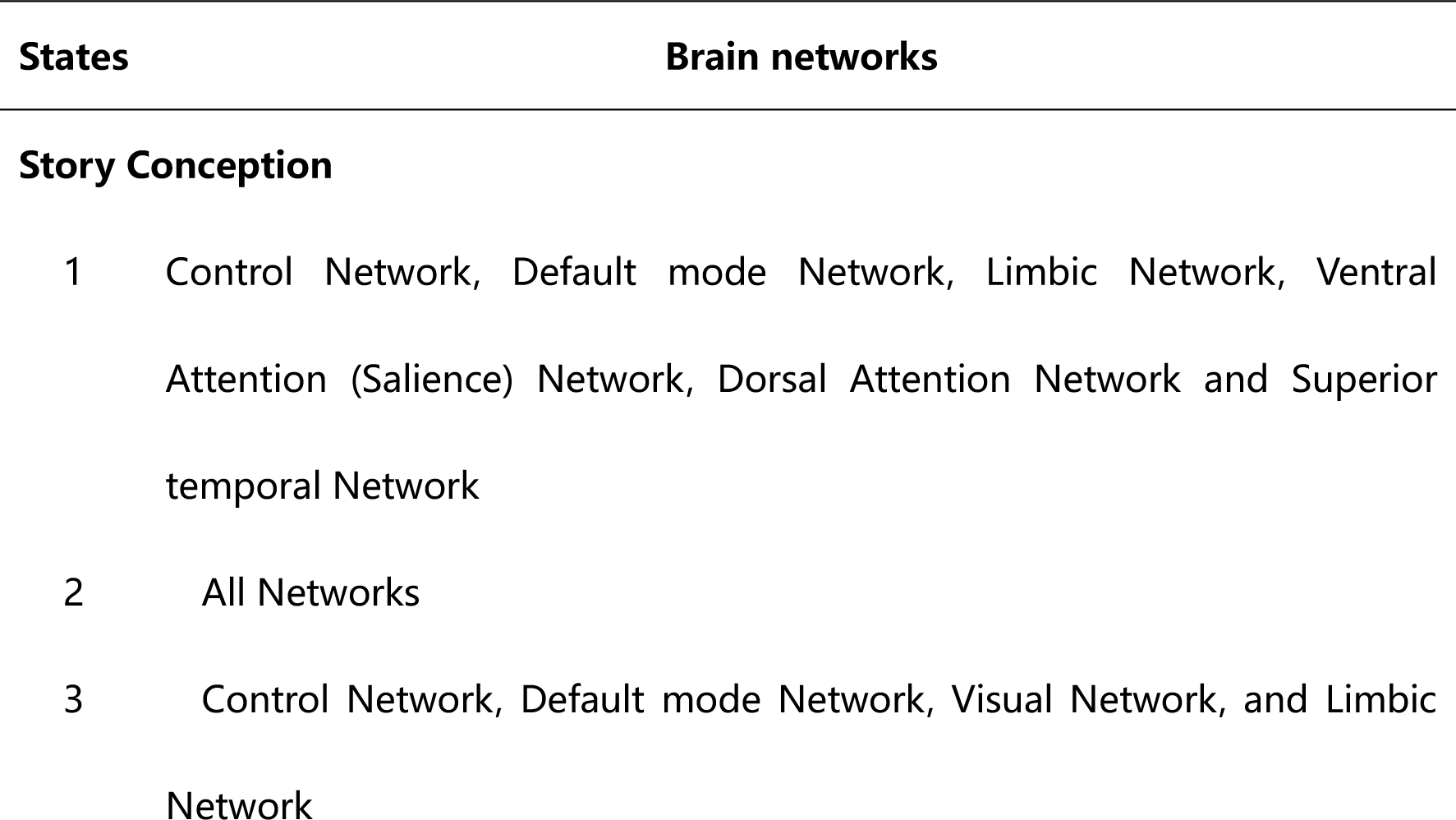
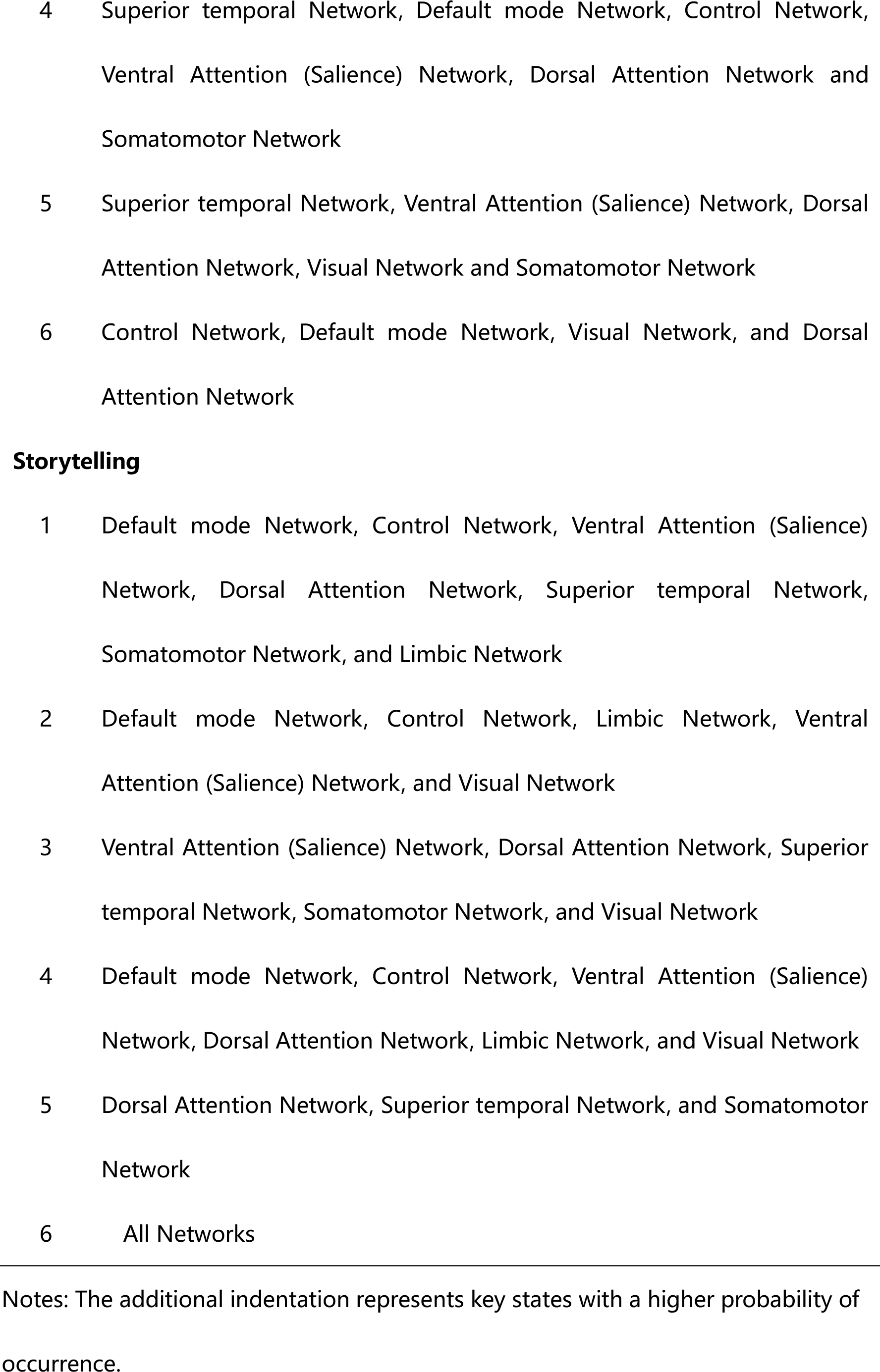
The synchronized brain networks in all states during the creative story conception and storytelling stages.

##### 3.3.1.2 The dwelling of states

In the creative story conception condition, the dwell time of state 2 and state 3 was significantly higher than that in the uncreative story conception condition (t=3.56, p=9.70×10^-4^ for state 2; t=5.85, p=7.58×10^-7^ for state 3). Meanwhile, the dwell time of state 6 was significantly higher in the creative than uncreative storytelling condition (t=4.96, p=1.35×10^-5^). Additionally, every participant’s average dwell time of each state exceeded zero, confirming that all participants have experienced each state.

##### 3.3.1.3 The transition pattern among key states

The number of state transitions was significantly lower in creative vs uncreative story conception (t=-5.46, p=2.70×10^-6^), as well as in creative vs uncreative storytelling (t=-4.38, p=8.23×10^-5^). Since only the story conception stage contained two key states, we only compared the number of transitions between state 2 and state 3 between creative and uncreative story conception. The results showed that the number of transitions between state 2 and state 3 was significantly higher under the creative than uncreative story conception condition (t=14.99, p=5.26×10^-8^).

Under creative story conception condition, we observed a significant increase in the probability of transitioning from state 2 to state 3 compared to uncreative conception condition. These specific changes are shown in **Figure 7A**. Similarly, during the creative storytelling stage, we found significant differences in state transition probabilities associated with state 6 compared to uncreative storytelling stage. These results are shown in **Figure 7B** and **Table 4**.

**Figure 7.**
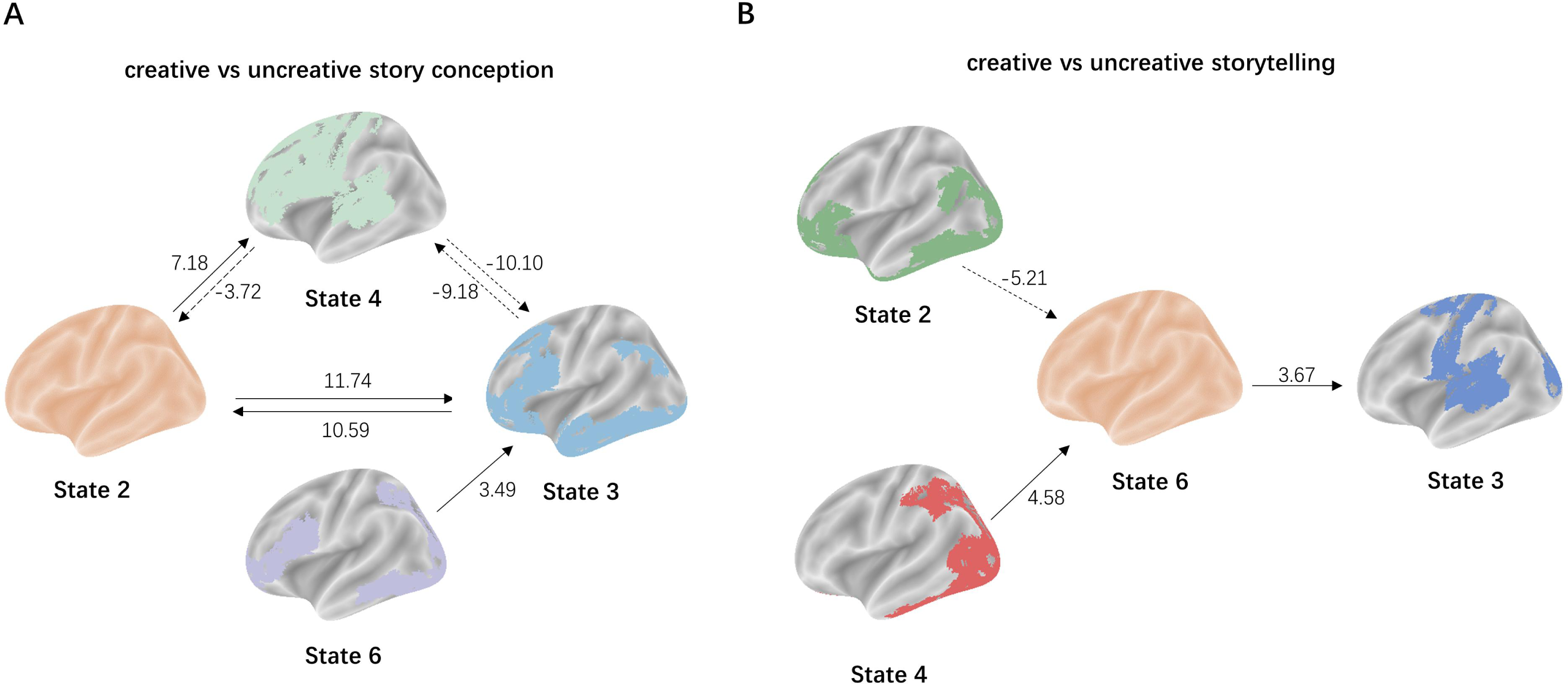
**A)** indicated the significant differences in the probability of transition during creative story conception compared to uncreative story conception, which were related to states 2 and 3. **B)** indicated the significant differences in the probability of transition during creative storytelling compared to uncreative story storytelling, which were related to states 6. The solid line represented a significant increase in the probability of state transition. The dashed line indicated a significant decrease in the probability of state transition. The numbers showed the t-values between creative and uncreative conditions in state transition probabilities.

**Table 4.**
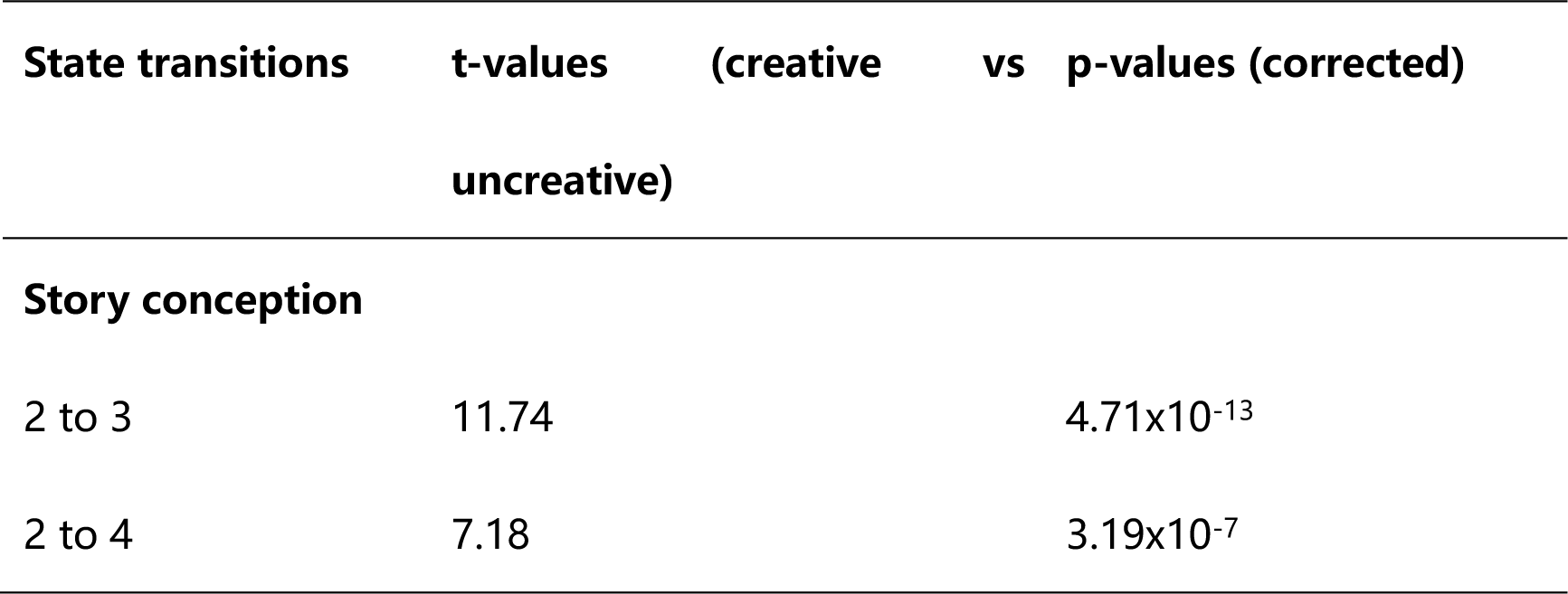
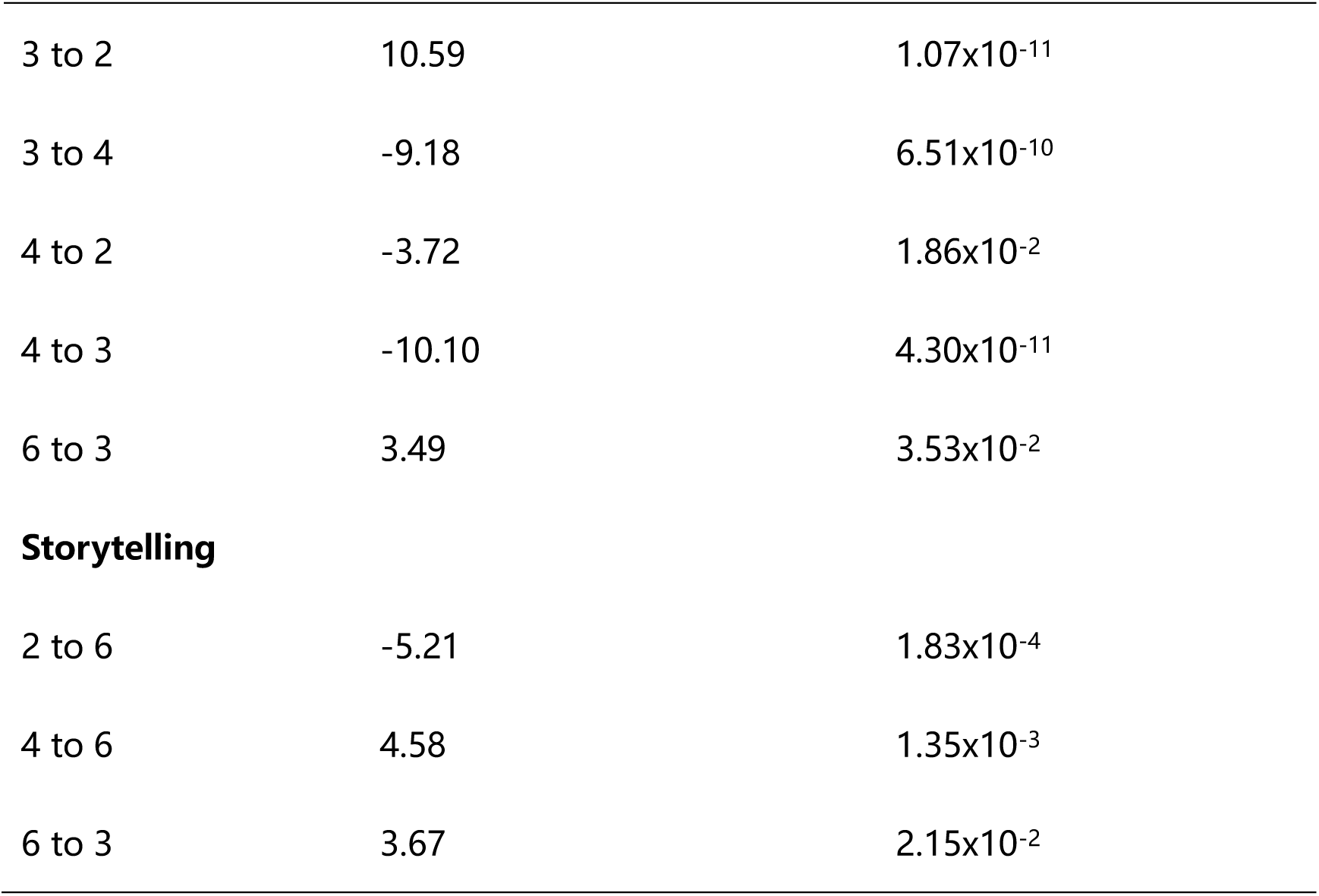
Significant differences between creative and uncreative conditions in state transition probabilities during the story conception and storytelling stage.

To present our results more intuitively, we created an animation of the state transitions of a participant during the 20 second creative story conception process, as shown in **example.mp4**.

##### 3.3.1.4 The time-varying pattern of key states

During the creative story conception period, the acceleration of the probability of state 2 occurrence was significantly less than 0 (mean=-5.60×10^-4^, SD=2.67×10^-3^, p=4.82×10^-6^), while the acceleration of the probability of state 3 occurrence was significantly greater than 0 (mean=4.77×10^-4^, SD=1.61×10^-3^, p=1.12×10^-10^). Similarly, during the uncreative story conception period, the acceleration of the probability of state 2 occurrence was also significantly less than 0 (mean=-5.30×10^-4^, SD=2.53×10^-3^, p=4.22×10^-6^), but there was no significant difference between the acceleration of the probability of state 3 occurrence and 0 (mean=1.07×10^-4^, SD=1.67×10^-3^, p=0.16).

In terms of storytelling, for both the creative and uncreative conditions, the acceleration of the probability of state 6 occurrence was significantly less than 0 (mean=-1.17×10^-4^, SD=2.36×10^-3^, p=2.88×10^-25^ for creative storytelling; mean=- 9.70×10^-4^, SD=2.27×10^-3^, p=1.12×10^-19^ for uncreative storytelling).

In order to provide a more intuitive description of our results, we plotted a mean quadratic curve fitting graph as an example, which represented the cumulative probabilities of state 2 and state 3 occurring over time for each participant in each trial during the creative story conception stage, as shown in the **Figure 8**.

**Figure 8.**
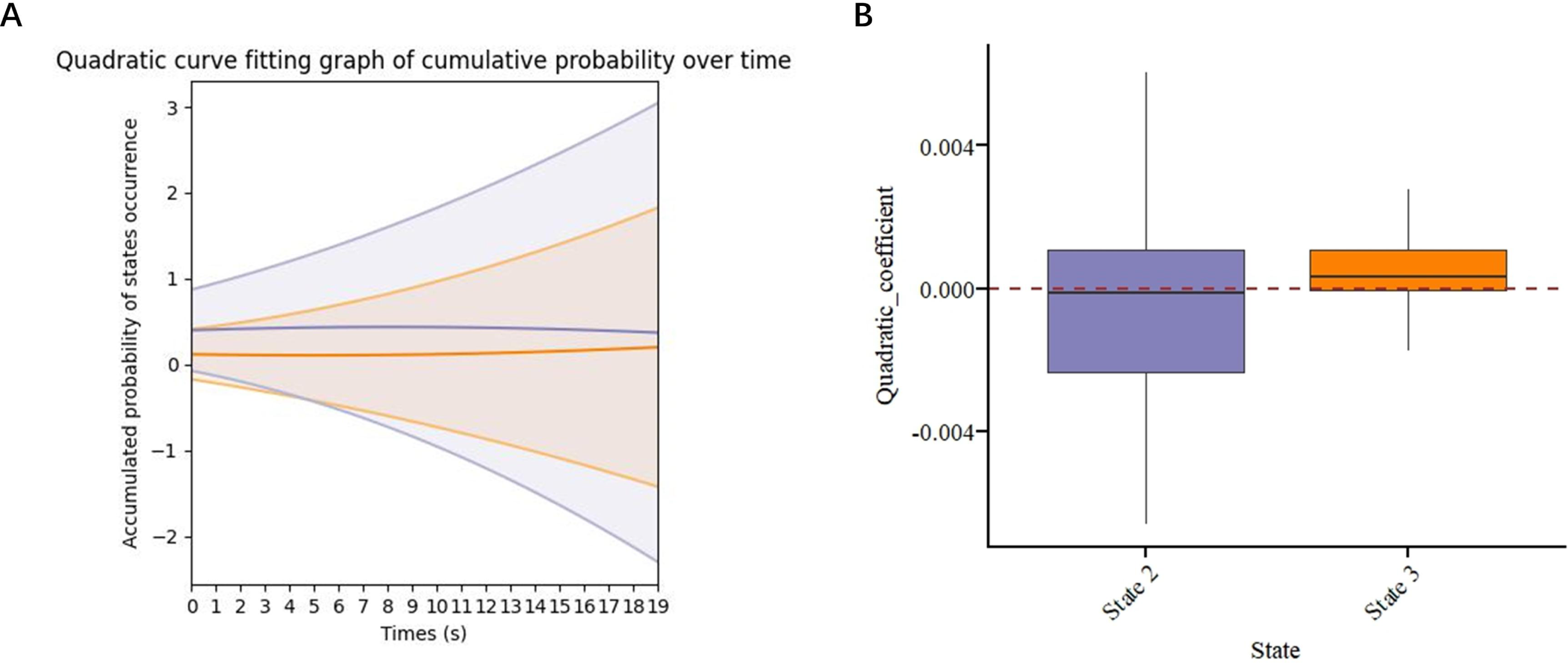
Shown the mean quadratic curve fitting graph and the quadratic coefficients of the cumulative probabilities of state 2 and state 3 occurring over time during the creative story conception phases. **A)** displayed quadratic curve fitting graph. The purple solid line represented the mean fitted curve of the cumulative probability of state 2 occurrence for each participant over time under each trail. The purple shaded area represented curves within one standard deviation range from the mean curve. The orange dashed line represented the mean fitted curve of the cumulative probability of state 3 occurrence for each participant over time under each trail. The orange shaded area represented curves within one standard deviation range from the mean curve. **B)** indicated the box chart of quadratic coefficients of fitted curve of the cumulative probability of state 2 and state 3 occurrence for each participant over time under each trail. The purple box represented state 2, and the orange box represented state 3.

**Figure 9.**
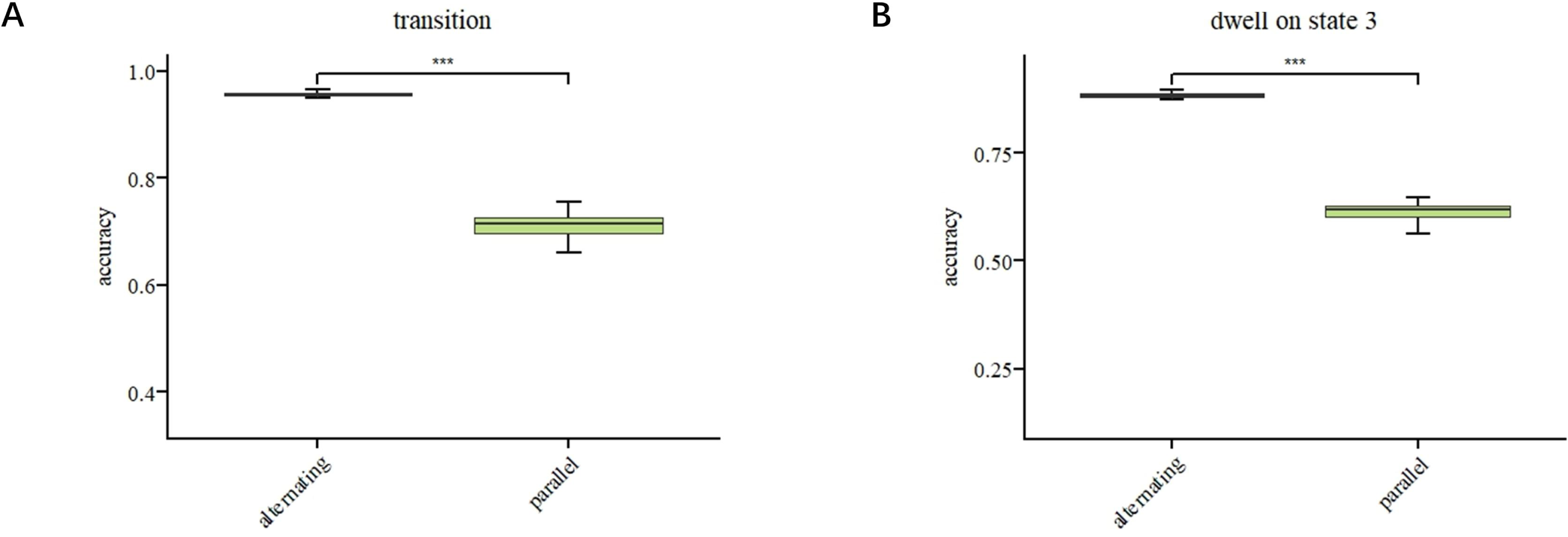
Shown the accuracy and differences in predicting times of transition between state 2 and state 3 **(A)**, and dwelling on state 3 **(B)**, using Alternating Co-attention model and Parallel Co-attention model, with changes in the number of hidden layers and epoch. The blue box represented the prediction accuracy of the Alternating Co-attention model, while the green box represented the prediction accuracy of the Parallel Co-attention model. The asterisk indicated that the p-value of the t-test was less than 0.001. The error bar indicates the range of one standard deviation from the mean prediction accuracy of the model, which can be positive or negative.

##### 3.3.1.5 The relationship between brain states and behavioral performance

The results of all regression analyses were shown in **Table 5**. RAT score was significantly correlated with the difference in time spent dwelling in state 2 (diff-state2) and state 3 (diff-state3) between creative and uncreative story conception, as well as with age. Using these three variables as independent variables and the RAT score as the dependent variable, we established a regression model. The results showed that the model was significant, with significant coefficients for diff-state3 between creative and uncreative story conception.

**Table 5.** Represented the results of regression analysis.

| Y | Adjusted<br>R<sup>2</sup> | F | p<br>(model) | Indicator | β | SE | t | p |
| --- | --- | --- | --- | --- | --- | --- | --- | --- |
| RAT score | 0.18 | 3.89 | 0.02 | diff- | 1.65 | 0.73 | 2.25 | 0.03 |
|  |  |  |  | state3 |  |  |  |  |
|  |  |  |  | age | -0.82 | 0.41 | -2.00 | 0.05 |
|  |  |  |  | diff- | 0.03 | 0.50 | 0.05 | 0.96 |
|  |  |  |  | state2 |  |  |  |  |
|  |  |  |  | Intercept | 36.84 | 8.62 | 4.27 | 0.0001 |
| **Creative writing score** | 0.18 | 5.53 | 0.008 | diff- | 0.97 | 0.39 | 2.51 | 0.02 |
|  |  |  |  | state3 |  |  |  |  |
|  |  |  |  | diff- | 0.24 | 0.23 | 1.00 | 0.32 |
|  |  |  |  | state2 |  |  |  |  |
|  |  |  |  | Intercept | 9.67 | 0.44 | 21.83 | 4.19x10<sup>-23</sup> |
| **Creative storytelling ( in scan ) score** | 0.41 | 14.76 | 1.81x10<sup>-5</sup> | diff- | 0.12 | 0.04 | 3.02 | 0.004 |
|  |  |  |  | state3 |  |  |  |  |
|  |  |  |  | Verbal IQ | 0.03 | 0.007 | 4.40 | 8.40x10<sup>-5</sup> |
|  |  |  |  | Intercept | -1.40 | 0.85 | -1.64 | 0.11 |

Notes: RAT = Remote Associates Test; diff-state2 = the difference in dwell time of state 2 between creative and uncreative story conception; diff-state3 = the difference in dwell time of state 3 between creative and uncreative story conception.

No independent variable significantly correlated with AUT score.

We found a significant correlation between Creative Writing scores, diff-state2 between creative and uncreative story conception and diff-state3 between creative and uncreative story conception. The regression model and coefficient for the diff-state3 between creative and uncreative story conception were both significant.

Finally, verbal IQ and diff-state3 between creative and uncreative story conception both correlated with scores on the in-scanner creative storytelling task. Regression results showed that both the model and the coefficients for the diff-state3 between creative and uncreative story conception were significant.

#### 3.3.2 Results about the parallel and alternating hypotheses of the dual process model

##### 3.3.2.1 The relationship between the dwells and transitions of states and brain networks involved in spontaneous and deliberate thinking

When dwelling in state 3, we observed that activation increased in brain networks such as the control network, ventral attention (salience) network, default mode network, and superior temporal network when compared to time points outside of state 3. Furthermore, at the point of transition between state 2 and state 3, the control network, dorsal attention network, ventral attention (salience) network, default mode network, and superior temporal network were all involved compared to the remaining time points.

##### 3.3.2.2 The Performance of Two Co-Attention Models

Results indicated that the Alternating Co-Attention model outperformed the Parallel Co-Attention model in terms of accuracy for predicting dwelling in state 3, and transitions between state 2 and state 3 (p=1.39×10^-32^; p=2.39×10^-47^). A box plot comparing the accuracy of the two models was presented in Figure 10.

### 3.4 Head motion considerations

To ensure that our results were not influenced by head motion, we calculated the mean framewise displacement (FD) for each participant. We found no significant correlation between FD and the dwell time of state 2 or state 3 during creative story conception, as well as the dwell time of state 6 during creative storytelling (r _state 2_=0.12; r _state 3_=0.03; r _state 6_=-0.09).

## 4. Discussion

In this study, we captured key brain states and dynamics during creative storytelling, examined their link to performance in creative tasks and explored mechanisms behind transitioning between and dwelling in brain states. Our findings firstly revealed that the probability of occurrence of a whole brain synchronization state during creative story conception and storytelling was significantly higher than under uncreative conditions. Additionally, the probability of occurrence of a synchronization state of the CDVL network was significantly higher during the creative story conception stage than during the uncreative conception stage. These states were identified as key states for creative thinking. Second, during the creative story conception period, the dwell life time of key states were significantly higher than during the uncreative story conception period. Similarly, the key state of the whole brain synchronization also had a longer dwell life time during the creative than the uncreative storytelling period. Third, the number of transitions and transition probability between key states were higher during creative than during uncreative story conception. Fourth, there were significant correlations between the difference of key states’ dwell life time and the creative behavior performance of the participants. Finally, compared to the Parallel Co-attention Model, the Alternating Co-attention Model performed better on predicting the dwell time points of key states and their transitions.

### 4.1 Linking the key brain states to idea generation and evaluation

We identified state 3, in which the brain synchronized across the CDVL network, as key state for creative thinking. Previous studies have found that the dwell life time of a state with increased resting-state functional connectivity between the control and default mode networks was related to creative performance (Beaty et al., 2018; Zhang et al., 2023). Additionally, in an fMRI study where participants were asked to design a book cover, researchers found higher activation of the control network, default mode network and limbic network during the idea evaluation stage compared to the idea generation stage (Ellamil et al., 2012). It is thus likely that state 3 reflects the evaluation stage of creative thinking.

Although previous creativity research has not reported on a state of whole brain synchronization, a LEiDA-based resting-state fMRI study on psilocybin suggested that such a state may reflect freely wandering thoughts, in this case under the influence of drugs (Lord et al., 2019). Other psilocybin studies reported increased whole brain communication, linked to reduced cognitive constraints and increased sensory associations (Petri et al., 2014; Roseman et al., 2014). Further, whole brain flexibility may serve as an important mediator between need for cognition and creative achievement (He et al., 2019). Given these findings, we suggest that this state of whole brain synchronization reflects the idea generation stage.

The temporal variations of the two key brain states throughout the entire creative thinking process further supported our speculations about the cognitive stages linking the two key states. Specifically, in both the creative story conception and storytelling stages, the growth rate of the probability density of whole brain synchronization slowed, while the that of synchronization across the CDVL network accelerated. Indeed, the pattern of temporal variations reflected their inherent dynamic characteristics, with idea generation being more prevalent in the early stages, and idea evaluation more prominent in the later stages. This finding also aligned with previous research using default mode network and control network information during different time periods to classify creative and uncreative AUT performances (Lloyd-Cox et al., 2022). Researchers found that the default mode network’s activity was more closely associated with classification accuracy in the initial stages compared to the middle and later stages. In contrast, the control network’s activity exhibited a weaker correlation with classification accuracy in the early stages than in the subsequent phases, corroborating the hypothesis of idea generation-evaluation within the creative thought process (Lloyd-Cox et al., 2022). Additionally, during the storytelling stage, the synchronized state of the whole brain exhibited a pattern of cumulative probability deceleration over time, which may be due to that participants gradually focus on oral production over time, possibly reducing idea generation.

Moreover, the correlation between brain states and creative performance links such states with different thinking stages. We found that the differences in the dwell life time of synchronization across the CDVL network between the creative and uncreative conditions could predict the performance of creative writing and convergent thinking. Importantly, although the state of whole brain synchronization was also associated with these two measures of creativity performance, synchronization across the CDVL network demonstrated a higher predictive effect. This could be attributed to the specific characteristics of the creative storytelling, creative writing and convergent thinking tasks. That is, idea evaluation is heavily needed to make a coherent story both in the creative storytelling and creative writing task. And previous studies have established a connection between the convergent thinking and idea evaluation (Cropley, 2006; Ulrich and Nielsen, 2020). Therefore, higher prediction performance of synchronization across the CDVL network might imply a potential link between this brain state and idea evaluation.

However, the precise meanings of these neural representations remain speculative. Under creative conditions, participants tend to use less frequent Chinese characters to form longer, more complex sentences. These basic textual feature differences might also contribute to neural differences. Nonetheless, by examining the network encompassing two key brain states, the temporal patterns of these states’ fluctuations, and their correlation with behavioral performances, we can preliminarily link the state of whole-brain synchronization to idea generation, and the state of synchronization across the CDVL network to idea evaluation.

### 4.2 Dynamic transition and circulation of idea generation and evaluation

Previous studies have hinted at a cyclical relationship between the generation and evaluation of ideas, yet definitive supporting evidence remained elusive. Building on this premise, we explored the transition patterns between brain states potentially linked to idea generation and evaluation. Our results demonstrated that during the creative conception stage, both the absolute number of transitions and the relative transition probability between the states of idea generation and evaluation were significantly higher than those observed in the uncreative conception stage. This increased frequency of mutual transition of the two key states supports the circulation between idea generation and evaluation proposed in the two-fold model of creativity (Kleinmintz et al., 2019). This dynamic feature serves as neural evidence for the cyclical nature of these two stages, offering a clearer exposition of the dynamic nature of brain network synchronization patterns specific to creative thinking and enriching our comprehension of the cognitive underpinnings of creative thinking.

However, the increase in the mutual transition between the two key states did not indicate a rise in overall state transition numbers. Our observations revealed that the key states potentially linked to idea generation and evaluation had longer dwell times under creative than uncreative conditions. This suggests that individuals may be more engaged in these two stages during the creative thinking process. This characteristic might enhance the stability of our brain, which refers to the consistency of brain activities over time. This is evidenced by a reduction in the total number of state transitions during the creative story conception and storytelling conditions compared to the uncreative conditions. These findings align with prior research showing reduced BOLD signal variation in creativity tasks (Roberts et al., 2020).

### 4.3 Possible alternating working pattern between spontaneous thinking and deliberate thinking

Our findings, in line with the dynamic framework of spontaneous thinking (Christoff et al., 2016), revealed that dwelling in state 3—brain synchronization across the CDVL network—was associated with fluctuations in large-scale networks, which indicated an amalgamation of both deliberate (control network) and spontaneous (comprising the ventral attention (salience) network, default mode network, and superior temporal network) thinking. State transitions between 2 and 3 were also tied to the fluctuations of large-scale networks, which could be interpreted as an integration of deliberate (control network) and spontaneous thinking processes (involving the dorsal attention network, ventral attention (salience) network, default mode network, and superior temporal network).

To test whether spontaneous and deliberate thinking alternate or run parallel during creative thinking, we used different co-attention mechanisms in deep learning to explore their interaction. We treated brain network activities related to spontaneous/deliberate thinking as two information sources and applied Parallel and Alternating Co-attention mechanisms to predict stable time points for brain state related with idea evaluation and to predict the time points of brain state transitions between key states. Our findings indicated that the Alternating Co-attention model outperformed the Parallel Co-attention model, suggesting that the creative thinking process was more consistent with the alternative model than the parallel model. However, our research is exploratory and may be influenced by various factors such as task characteristics. Therefore, it does not entirely negate the parallel hypothesis. Additionally, a new working model has been proposed, which posits that competing intuitions (spontaneous thinking) determine the involvement of deliberate thinking. Specifically, when the conflict between intuitions reached a threshold, deliberate thinking took over until the conflict subsided (De Neys, 2022). Our results may also align with this model. Nevertheless, our findings provide initial support for the alternating hypothesis and offer methods and insights for future studies.

## 5. Conclusion

Using an approach capable of capturing brain dynamics, we delineated the ongoing dynamic cycles between the idea generation and evaluation stages during creative storytelling, providing neurobiological evidence for the circulation between these two stages proposed by the two-fold model. By employing deep learning techniques, we determined that spontaneous and deliberate thinking drive the transition between and maintenance of the two stages through an alternating mode that is difficult to directly observe. These findings enhance our comprehension of creative thinking, and serve to settle the debate of how the two stages operate and interact. More generally, our approaches from dynamic and algorithmic standpoints demonstrate potentials for understanding cognitive mechanisms underlying human thinking.

## Acknowledgments

This research was supported by the STI 2030—Major Projects (2021ZD0200500), the National Natural Science Foundation of China (31970977, 31571155), the Interdisciplinary Research Funds of Beijing Normal University and the Fundamental Research Funds for the Central Universities (2015KJJCB28).

**example.** Displayed the brain state transition process per second of a participant during a 20 second creative story conception process.

## Highlights

- A state with whole brain synchronization is related to idea generation
- Synchronization of 4 networks including DMN and control connects to idea evaluation
- Idea generation and evaluation circulate during creative story conception
- Spontaneous and deliberate thinking works alternately during creative storytelling

## Conflict of interest statement

The authors declare that they have no known competing financial interests or personal relationships that could have appeared to influence the work reported in this paper.

## Data and Code Availability Statement

Some or all data, models, or code generated or used during the study are available from the corresponding author, Liu, upon reasonable request.

## Supplementary Materials

**Figure S1.**
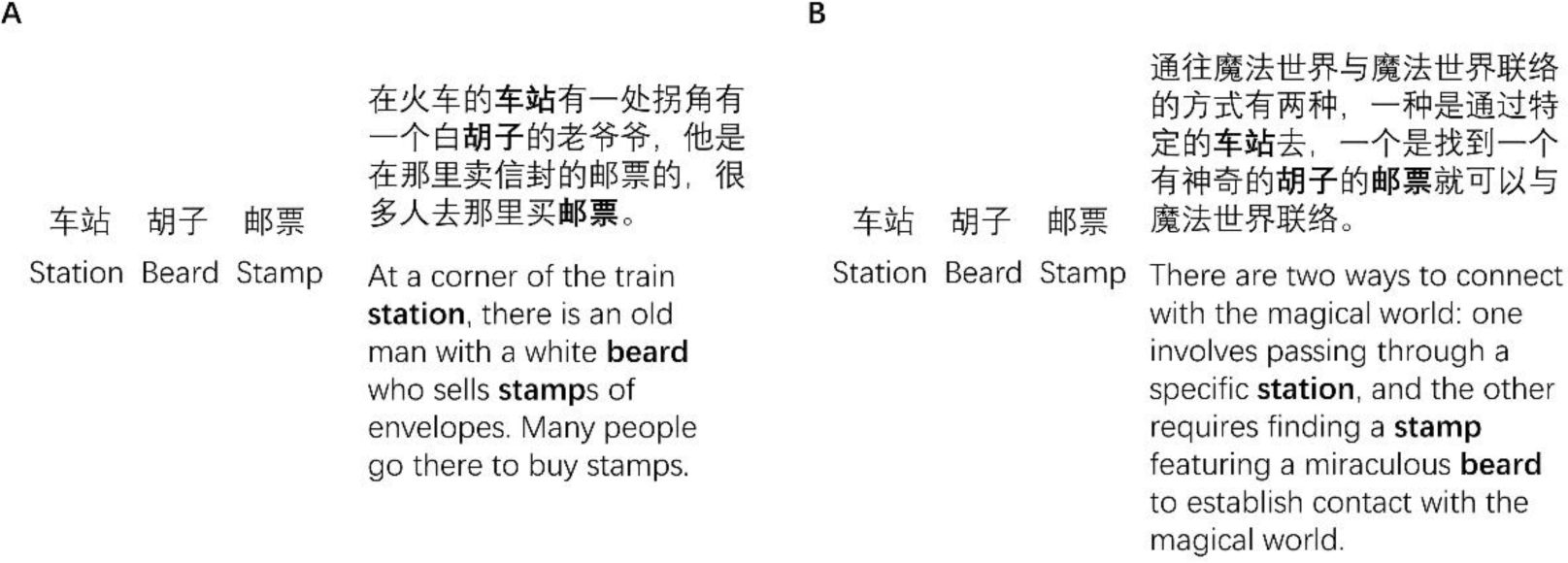
Two stories told by different participants during the scanning process based on the same words. **A)** belonged to the uncreative condition. **B)** belonged to the creative condition. The left column lists three key words, while the right column presents a story prompted by these words, with the keywords bolded for emphasis.

**Figure S2.**
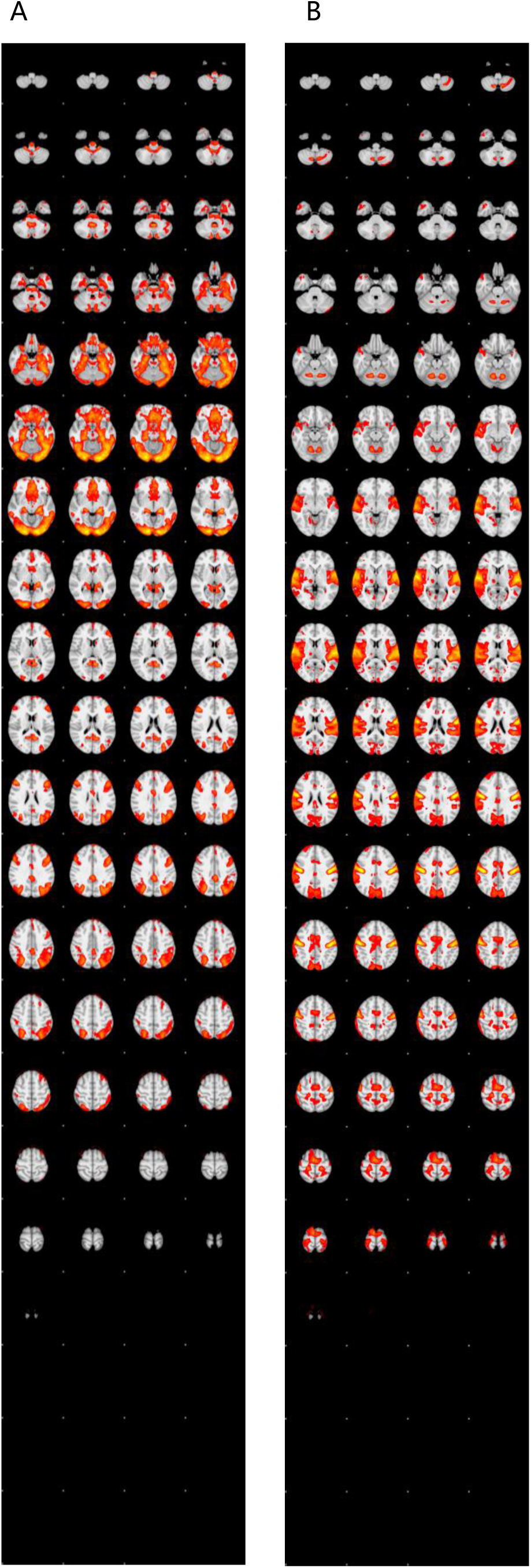
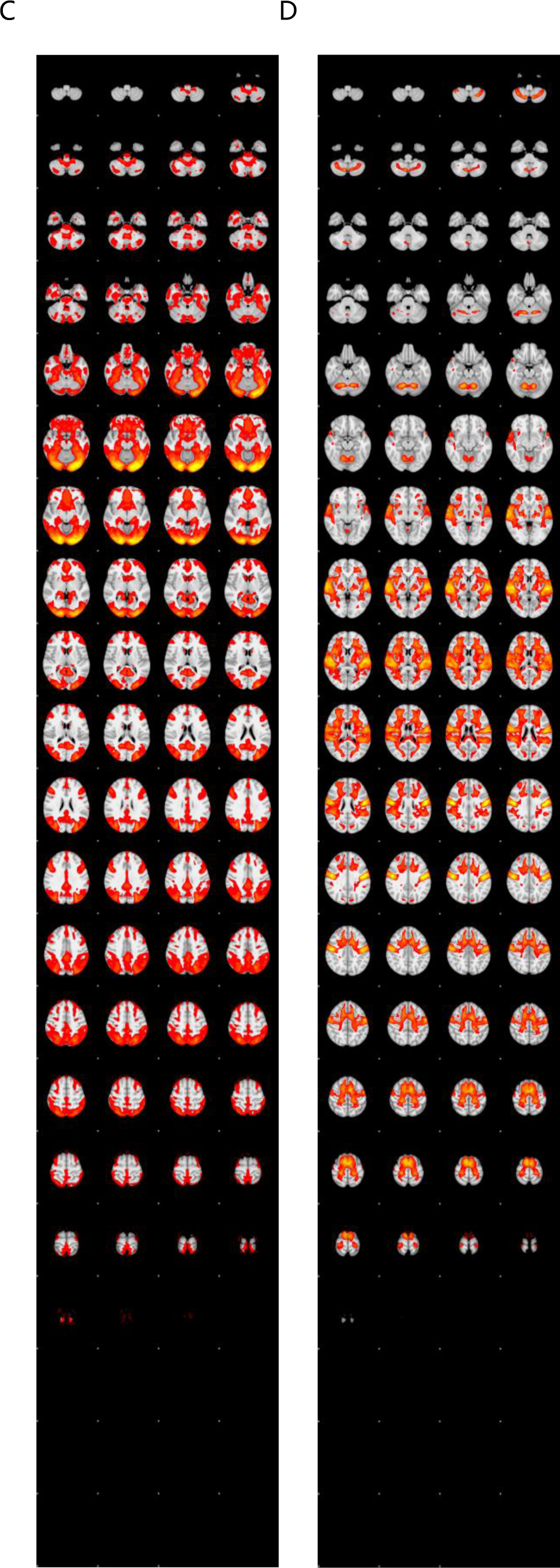
Displayed whole brain activation maps under different contrasts. A) The results of contrast: creative conception-rest. B) The results of contrast: uncreative conception-rest. C) The results of contrast: creative storytelling-rest. D) The results of contrast: uncreative storytelling-rest.

**Table S1.**
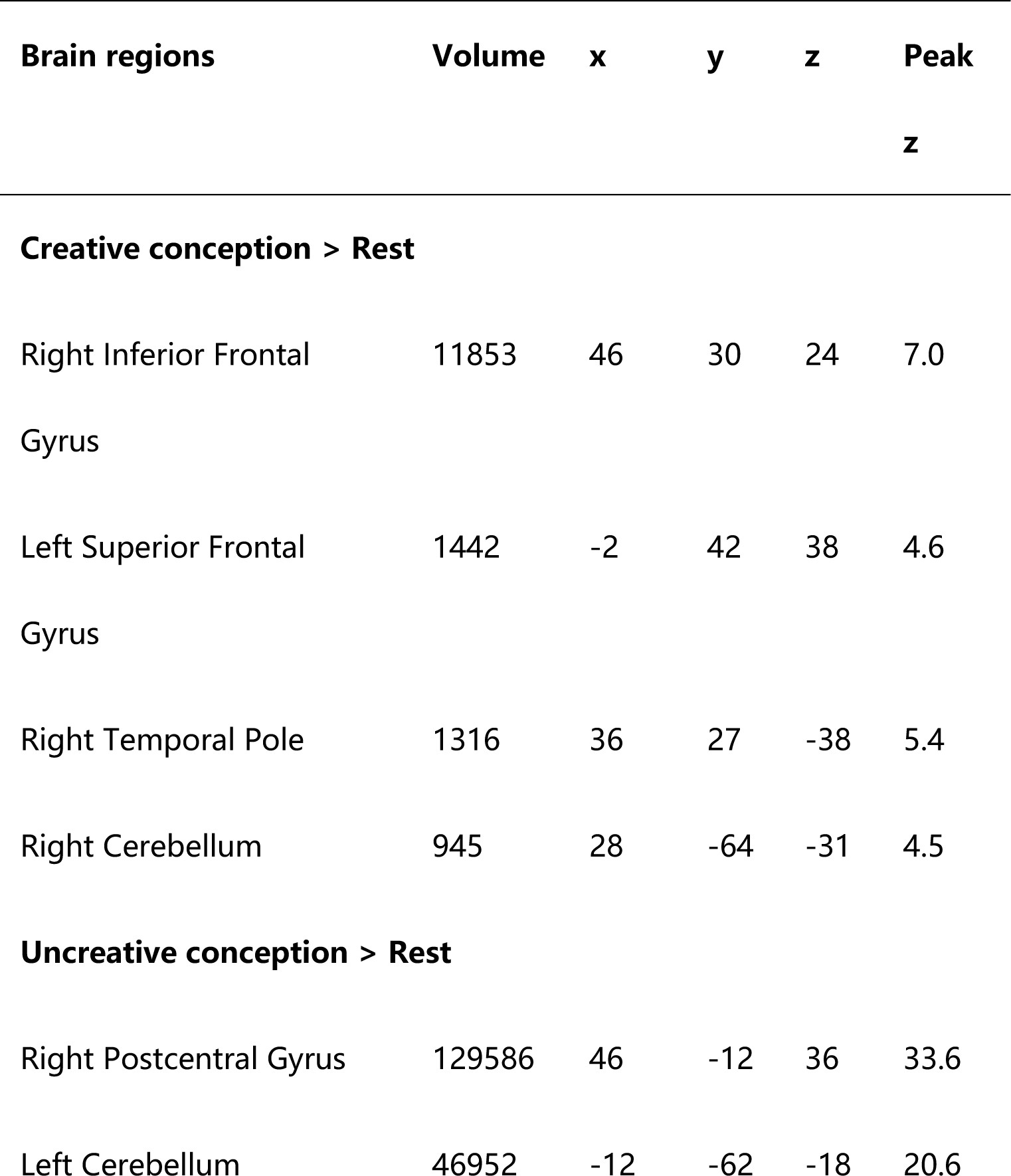
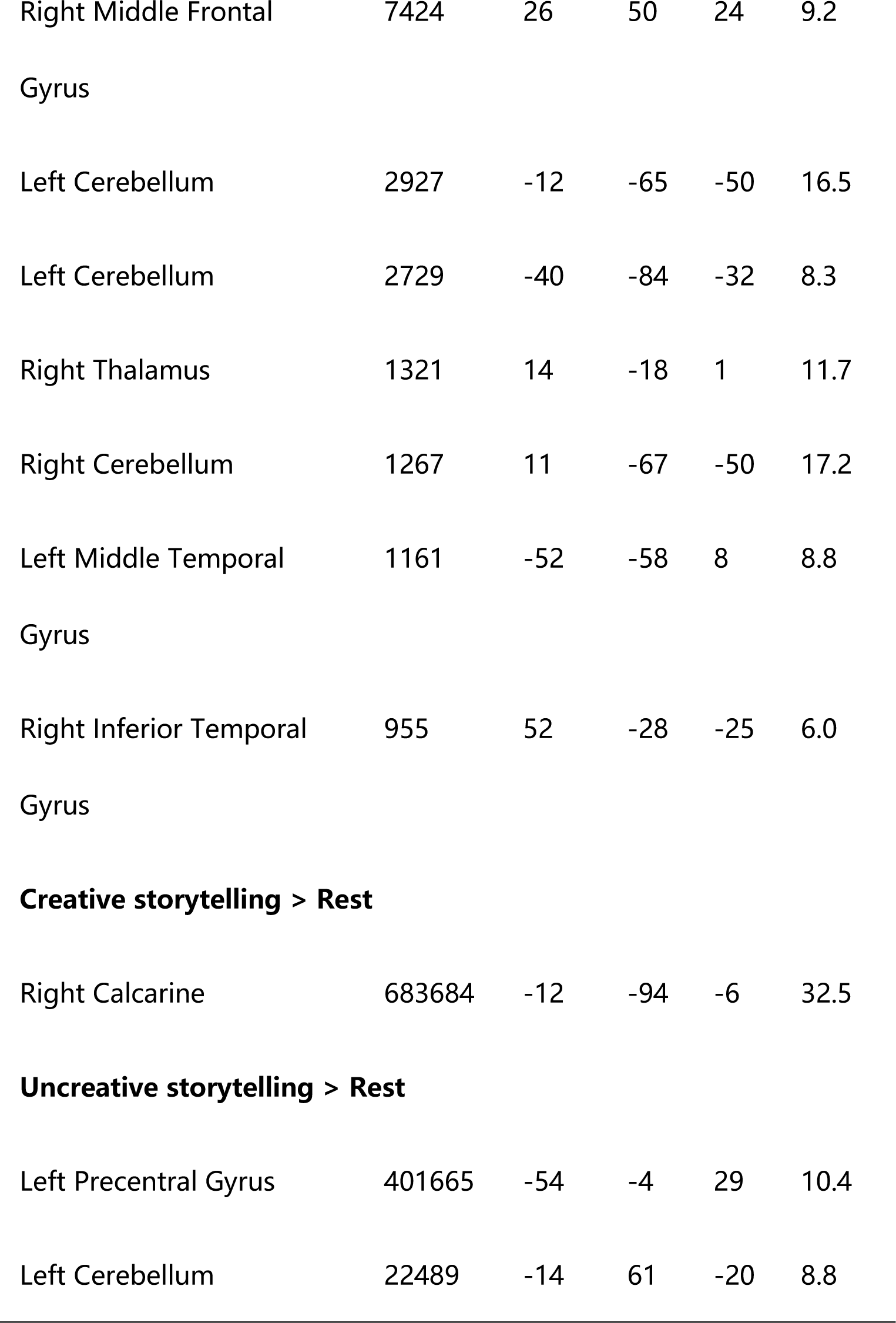
Whole brain activation results under different contrasts.

| Brain regions | Volume | x | y | z | Peak<br>z |
| --- | --- | --- | --- | --- | --- |
| **Creative conception > Rest** |  |  |  |  |  |
| Right Inferior Frontal Gyrus | 11853 | 46 | 30 | 24 | 7.0 |
| Left Superior Frontal Gyrus | 1442 | -2 | 42 | 38 | 4.6 |
| Right Temporal Pole | 1316 | 36 | 27 | -38 | 5.4 |
| Right Cerebellum | 945 | 28 | -64 | -31 | 4.5 |
| **Uncreative conception > Rest** |  |  |  |  |  |
| Right Postcentral Gyrus | 129586 | 46 | -12 | 36 | 33.6 |
| Left Cerebellum | 46952 | -12 | -62 | -18 | 20.6 |
| Right Middle Frontal<br>Gyrus | 7424 | 26 | 50 | 24 | 9.2 |
| Left Cerebellum | 2927 | -12 | -65 | -50 | 16.5 |
| Left Cerebellum | 2729 | -40 | -84 | -32 | 8.3 |
| Right Thalamus | 1321 | 14 | -18 | 1 | 11.7 |
| Right Cerebellum | 1267 | 11 | -67 | -50 | 17.2 |
| Left Middle Temporal<br>Gyrus | 1161 | -52 | -58 | 8 | 8.8 |
| Right Inferior Temporal<br>Gyrus | 955 | 52 | -28 | -25 | 6.0 |
| **Creative storytelling > Rest** |  |  |  |  |  |
| Right Calcarine | 683684 | -12 | -94 | -6 | 32.5 |
| **Uncreative storytelling > Rest** |  |  |  |  |  |
| Left Precentral Gyrus | 401665 | -54 | -4 | 29 | 10.4 |
| Left Cerebellum | 22489 | -14 | 61 | -20 | 8.8 |

## Notes

### Competing Interest Statement

The authors have declared no competing interest.

